# Exploring the Effects of Teas with Different Fermentation Levels and Black Coffee on the Body via the Urine Proteome

**DOI:** 10.1101/2025.07.02.662780

**Authors:** Yuzhen Chen, Youhe Gao

## Abstract

Tea and black coffee, two of the most widely consumed beverages worldwide, play important roles in supporting overall health. Urinary proteins do not originate directly from the beverages. Instead, changes in the urine proteome reflect the changes in the body influenced by beverage consumption, rather than their metabolites. Can the effects of teas with different fermentation levels and black coffee on the body be explored via the urine proteome? In this study, urine samples were collected from rats before and after seven consecutive days of consuming green tea, oolong tea, black tea, Pu-erh tea, or black coffee. Both before-and-after comparisons and between-group comparisons were performed, and the samples were analyzed using liquid chromatography coupled with tandem mass spectrometry. The results showed that the urine proteome reflected the changes in rats after one week of consuming teas with different fermentation levels or black coffee. Biological processes and pathways enriched from differential proteins included fat cell differentiation, lipid metabolism, glucose metabolism, fatty acid transport, and immune response. Furthermore, the effects of teas with different fermentation levels and black coffee on the body exhibited a high degree of specificity. Additionally, several differential proteins identified in this study have been reported as biomarkers for diseases such as cancer and cardiovascular diseases. This suggests that beverage consumption, including tea and black coffee, should be considered in clinical application of urine biomarkers of diseases and related studies. And the use of biomarker panels may be necessary to improve accuracy. In conclusion, the urine proteome provides a comprehensive and systematic reflection of the overall effects of all components in tea and black coffee on the body, and can distinguish changes in the body after consuming teas with different fermentation levels and black coffee.

## 1 Introduction

Tea is the second most consumed beverage in the world after water and contains a variety of bioactive components, including polyphenols, theaflavins, thearubigins, and caffeine, which possess antiviral, antioxidant, and anti-inflammatory properties^1^. Numerous studies have shown that tea has beneficial effects in reducing the risk of cardiovascular disease, stroke^2^, type 2 diabetes mellitus^3^, hypertension^4^, various cancers^5^, and dementia^6^, as well as improving mood^7^, anti-aging^8^, and anti-obesity^9^. Tea can be classified based on the levels of fermentation into green tea (unfermented), oolong tea (partially fermented), black tea (completely fermented), and Pu-erh tea (drastically fermented and aged)^10^. The level of fermentation affects the content of bioactive components in the tea^11^.

Coffee is also one of the most widely consumed beverages worldwide. It contains more than a thousand compounds, including caffeine, chlorogenic acid, diterpenes, and trigonelline^12^. Coffee plays a positive role in reducing the mortality of cardiovascular diseases and the incidence of stroke^13^, lowering the risk of cancer^14^, Parkinson’s disease^15^, and type 2 diabetes mellitus^16^, as well as anti-obesity^17^.

Proteomics research reveals the composition and dynamics of proteins within cells or organisms by analyzing protein structure, expression, post-translational modifications, and protein interactions^18^. Urine, not strictly regulated by homeostatic mechanisms, can accommodate and accumulate more changes, reflecting changes in all organs and systems of the body earlier and more sensitively^19^. Tea and black coffee also contain unidentified components that may contribute to overall health. Since urinary proteins are basically not derived directly from these beverages, changes in the urine proteome reflect the changes in the body under the influence of beverage rather than the presence of beverage metabolites themselves. Thus, the use of the urine proteome offers a comprehensive and systematic approach to exploring the overall effects of all the components of these beverages.

Various factors, such as age, genetics, gender, diet, and exercise, inevitably affect the urine proteome. Therefore, it is crucial to reduce the interference of extraneous factors in experiments. Animal models, whose genetic and environmental factors can be controlled, are suitable choices^20^.

Can we leverage urine’s ability to comprehensively, systematically, and sensitively reflect the body’s state to meticulously explore the effects of widely consumed tea and black coffee, and seek the differences among teas with different fermentation levels and black coffee? In this study, we selected four types of tea with different fermentation levels (commercial green tea, oolong tea, black tea, and Pu-erh tea) and commercial black coffee. The effects and differences of teas and black coffee on rats were explored via the urine proteome, aiming to provide new clues for the study of their actions. Meanwhile, biomarkers affected by teas and black coffee were explored, so as to provide references for the clinical application of urine biomarkers of diseases and related studies (Figure 1).

**Figure 1.**
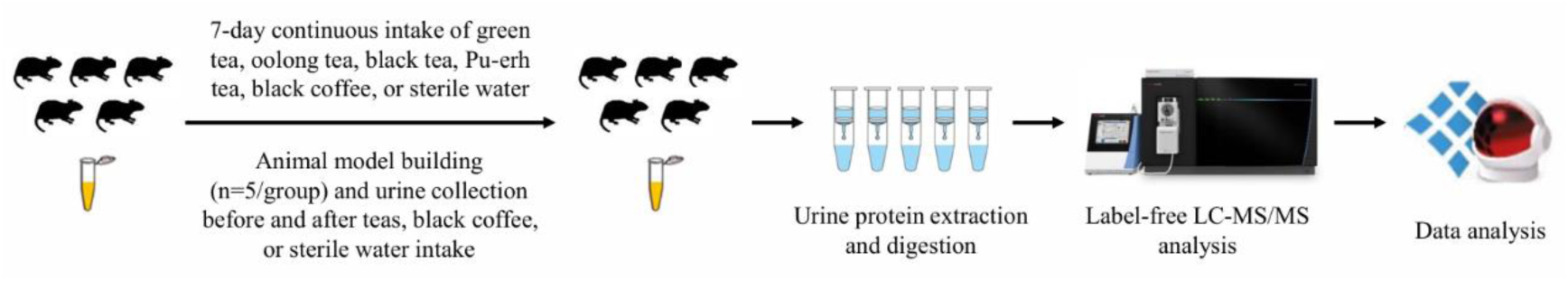
Technical route for exploring the effects of teas with different fermentation levels and black coffee on the body via the urine proteome.

## 2 Materials and Methods

### 2.1 Urine Collection

Thirty healthy male Sprague Dawley (SD) rats (200 ± 20 g) aged 6–7 weeks were purchased from Beijing Vital River Laboratory Animal Technology Co., Ltd, Beijing, China. All rats were kept under standard conditions [room temperature (22±1) ℃, humidity 65%-70%]. Animal experiments were reviewed and approved by the Ethics Committee of the College of Life Sciences, Beijing Normal University (No. CLS-AWEC-B-2022-003).

The thirty rats were randomly assigned into six groups (five rats per group). After a 3-day acclimation period under standard conditions, the rats were uniformly placed in metabolic cages to collect urine samples for 12 h. The experimental groups consumed commercial green tea, oolong tea, black tea, Pu-erh tea, or black coffee, while the control group consumed sterile water. The Pu-erh tea was prepared by dissolving 96.5 mg of tea powder in 500 mL of sterile water. The black coffee was prepared by diluting 33 mL of coffee concentrate in 547 mL of sterile water. All the beverages and sterile water were placed in 250 mL drinking bottles, and the rats were allowed to consume food and drink ad libitum. The teas, black coffee, and sterile water were replaced with new ones every day. After 7 days of treatment, rats were uniformly placed in metabolic cages to collect urine samples for 12 h. The urine samples were then stored at −80℃.

### 2.2 Urine Sample Preparation for Label-free Analysis

The collected urine samples were centrifuged at 12 000 ×*g* for 40 min at 4℃. The supernatant was transferred to a new centrifuge tube. Pre-cooled anhydrous ethanol, three times the volume of the supernatant, was added, mixed thoroughly, and then precipitated overnight at −20℃. Following centrifugation at 12 000 ×*g* for 30 min at 4℃, the supernatant was discarded. The protein precipitate was then suspended in an appropriate lysis buffer (8 mol/L urea, 2 mol/L thiourea, 25 mmol/L dithiothreitol, and 50 mmol/L Tris). After being fully dissolved, the sample was centrifuged again at 12 000 ×*g* for 30 min at 4℃. The supernatant was transferred to the new centrifuge tube to obtain urine protein extract. Protein concentration was quantified using the Bradford kit assay (Applygen, Beijing, China).

Using filter-aided sample preparation (FASP) method^21^, a total of 100 μg of protein was added to a 1.5 mL centrifuge tube. 25 mmol/L NH_4_HCO_3_ solution was added to make a total volume of 200 μL. 20 mmol/L dithiothreitol solution (DTT, Sigma) was added, vortexed, and mixed well. The sample was heated in a metal bath at 97℃ for 10 min. After cooling to room temperature, 50 mmol/L iodoacetamide solution (IAA, Sigma) was added, vortexed, and mixed well, and reacted at room temperature without light for 40 min. 200 μL of UA solution (8 mol/L urea, 0.1 mol/L Tris-HCl, pH 8.5) was added to a 10 kD ultrafiltration tube (Pall, Port Washington, NY, USA) and centrifuged twice at 14 000 ×*g* for 5 min at 18℃. The treated protein sample was added and centrifuged at 14 000 ×*g* for 40 min at 18℃. 200 μL of UA solution was then added and centrifuged at 14 000 ×*g* for 40 min at 18℃, repeated once. 25 mmol/L NH_4_HCO_3_ solution was added and centrifuged at 14 000 ×*g* for 40 min at 18℃, repeated once. The samples were digested overnight at 37℃ with trypsin (Trypsin Gold, Promega, Madison, WI, USA) at an enzyme-to-protein ratio of 1∶50. The digested peptides were eluted from ultrafiltration membranes, desalted with HLB columns (Waters, Milford, MA), dried in a vacuum desiccator, and stored at −80℃.

### 2.3 Liquid Chromatography Coupled with Tandem Mass Spectrometry Analysis

The digested peptides were dissolved in 0.1% formic acid, and the peptide concentration was quantified using a BCA kit. The peptides were then diluted to a final concentration of 0.5 μg/μL. A mixed peptide sample was prepared by combining 3.3 μL from each sample, and separation was performed using a high pH reversed-phase peptide fractionation kit (Thermo Fisher Scientific, Rockford, IL, USA). Ten fractions were collected by centrifugation, dried in a vacuum desiccator, and redissolved in 0.1% formic acid. The indexed retention time (iRT) reagent (Biognosis, Schlieren, Switzerland) was then added at a ratio of 1:10 (iRT-to-sample).

Ten fractions obtained from the high pH reversed-phase peptide fractionation kit were analyzed by mass spectrometry in a data-dependent acquisition (DDA) mode to generate a spectral library. 1 μg of the peptide from each sample was loaded onto a reversed-phase C18 trap column (75 μm × 2 cm, 3 μm) at a flow rate of 0.3 μL/min and separated with a reversed-phase analytical column (50 µm × 15 cm, 2 µm). A 90-minute gradient elution was applied using mobile phase A (0.1% formic acid) and mobile phase B (0.1% formic acid in 80% acetonitrile) (Table 1).

**Table 1.** 90-min gradient elution procedure.

| Time | Duration | Flow (nL/min) | %B |
| --- | --- | --- | --- |
| 00:00 | 00:00 | 300 | 4 |
| 02:00 | 02:00 | 300 | 6 |
| 62:00 | 60:00 | 300 | 22 |
| 78:00 | 16:00 | 300 | 35 |
| 80:00 | 02:00 | 300 | 90 |
| 90:00 | 10:00 | 300 | 90 |

Analysis was performed using an Orbitrap Fusion Lumos Tribrid Mass Spectrometer (Thermo Fisher Scientific, Waltham, MA, USA). The spray voltage was set to 2.25 kV. A full MS scan was acquired within a 350-1 550 *m/z* range with a resolution of 120 000. The MS/MS scan was acquired in Orbitrap mode with a resolution of 30 000. The HCD collision energy was set to 30%. The top 20 precursors were selected and subjected to a 30-second dynamic exclusion period. The DDA results were then imported into Proteome Discoverer software (version 2.1, Thermo Fisher Scientific) for searching against the *Rattus norvegicus* database using SEQUEST HT. The search results were used to establish the DIA method. The width and number of windows were calculated based on the *m/z* distribution density.

Individual samples were analyzed using the data-independent acquisition (DIA) mode, with each sample run in triplicate. The LC settings for DIA mode were identical to those used in DDA mode. The spray voltage was set to 2.3 kV. A full MS scan was acquired within a 350-1 500 *m/z* range with a resolution of 60 000. The MS/MS scan was acquired in Orbitrap mode in the range of 200-2 000 *m/z* with a resolution of 30 000. The HCD collision energy was set to 32%. After every 9-10 runs, a single DIA analysis of the pooled peptides was performed to control the quality of the whole analytical process.

### 2.4 Database Searching and Data Processing

The results were then imported into Spectronaut Pulsar (version 19, Biognosys AG, Schlieren, Switzerland) software for analysis and processing. The peptide intensity was calculated by summing the peak areas of the respective fragment ions for MS^2^, while protein intensity was calculated by summing the peptide intensities.

### 2.5 Data Analysis

In this study, both before-and-after comparisons and between-group comparisons were performed. To minimize the effects of individual differences, differential proteins were screened by comparing urinary proteins before and after teas or black coffee consumption. The screening criteria were as follows: fold change (FC) ≥1.5 or ≤ 0.67, and *p* < 0.05 by two-tailed paired *t*-test analysis. To avoid the effects of short-term growth and development of rats, urinary proteins of the experimental and control groups were compared after rats consumed teas, black coffee, or sterile water. Differential proteins were screened with the following criteria: FC ≥1.5 or ≤ 0.67, and *p* < 0.05 by two-tailed unpaired *t*-test analysis.

Functional enrichment analysis of differential proteins was performed using the UniProt database (https://www.uniprot.org/) and the Database for Annotation, Visualization, and Integrated Discovery (DAVID) (https://davidbioinformatics.nih.gov/). Functional analysis was further supported by searching the PubMed database (https://pubmed.ncbi.nlm.nih.gov/) for relevant studies in the literature.

## 3 Results

In this study, rats showed no obvious preference for any of the teas with different fermentation levels or black coffee. Sixty samples from both the experimental and control groups were analyzed by liquid chromatography coupled with tandem mass spectrometry. A total of 1 647 proteins were identified based on the criteria that each protein contained at least two specific peptides and a false discovery rate (FDR) <1% at the protein level.

### 3.1 Comparative Analysis Before and After Consuming Teas and Black Coffee

#### 3.1.1 Comparison Before and After Green Tea Consumption

By comparing the urinary proteins of rats before and after green tea consumption, a total of 61 differential proteins were identified under the screening conditions of FC ≥ 1.5 or ≤ 0.67 and *p* < 0.05 by two-tailed paired *t*-test analysis. Detailed information on these differential proteins is listed in Table S1. To assess the possibility of random generation of the identified differential proteins, a randomized grouping test was performed on the total proteins. Ten samples before and after green tea consumption were randomly divided into two new groups, resulting in a total of 126 combinations. These combinations were then screened for differences based on the same criteria (FC ≥ 1.5 or ≤ 0.67, *p* < 0.05). The average number of differential proteins yielded was 39.84, indicating that at least 34.69% of the differential proteins were not randomly generated.

##### 3.1.1.1 Analysis of Differential Proteins Identified in the Green Tea Group

Among the differential proteins identified before and after green tea consumption, CCN family member 1 and Cadherin 26 exhibited changes from presence to absence. This means that they were identified in the urinary samples of rats before green tea consumption but not in those after consumption. CCN family member 1 (FC = 0, *p* = 1.65×10^-03^) possesses the function of growth factor binding, participates in the regulation of multicellular organismal process. It has been reported as an early marker for infarct size and left ventricular dysfunction in patients with ST-segment elevation myocardial infarction^22^. Cadherin 26 (FC = 0, *p* = 1.43×10^-02^) is involved in biological processes including the adherens junction organization, calcium-dependent cell-cell adhesion via plasma membrane cell adhesion molecules, CD4-positive alpha-beta T cell activation, cell migration, and cell morphogenesis.

45 kDa calcium-binding protein (Stromal cell-derived factor 4) (FC = 7.90, *p* = 6.76×10^-04^) exhibited the smallest *p*-value and the second-highest FC value. It possesses the function of calcium ion binding and participates in fat cell differentiation. Studies have shown that SDF4 expression is elevated in various cancer cell types, especially those with high proliferative and metastatic potential. Serum SDF4 levels have been proposed as a diagnostic biomarker for malignancies such as gastric cancer^23^. In addition, SDF4 has been reported as a potential biomarker for pancreatic cancer^24^. SDF4 may also serve as a potential biomarker and therapeutic target for glioblastoma^25^. N-acetylneuraminate synthase (FC = 0.16, *p* = 4.72×10^-02^) exhibited the third-smallest FC value. It possesses synthase activity, and is involved in carbohydrate biosynthetic process, CMP-N-acetylneuraminate biosynthetic process, glycosylation, and N-acetylneuraminate biosynthetic process. It has also been reported as a potential prognostic biomarker for aggressive prostate cancer^26^.

Tissue-type plasminogen activator (FC = 0.16, *p* = 4.01×10^-02^) exhibited the fourth-smallest FC value. It is involved in biological processes including the regulation of synaptic plasticity, glutamatergic synaptic transmission, trans-synaptic signaling by BDNF, and responses to cAMP, fatty acid, and hypoxia.

##### 3.1.1.2 Analysis of Biological Pathways Enriched from Differential Proteins in the Green Tea Group

Biological process enrichment analysis of the identified differential proteins was performed using the DAVID database (Figure 2). These differential proteins were mainly involved in biological processes such as homophilic cell adhesion via plasma membrane adhesion molecules, negative regulation of oxidative stress-induced intrinsic apoptotic signaling pathway, fat cell differentiation, UV protection, and pulmonary valve morphogenesis.

**Figure 2.**
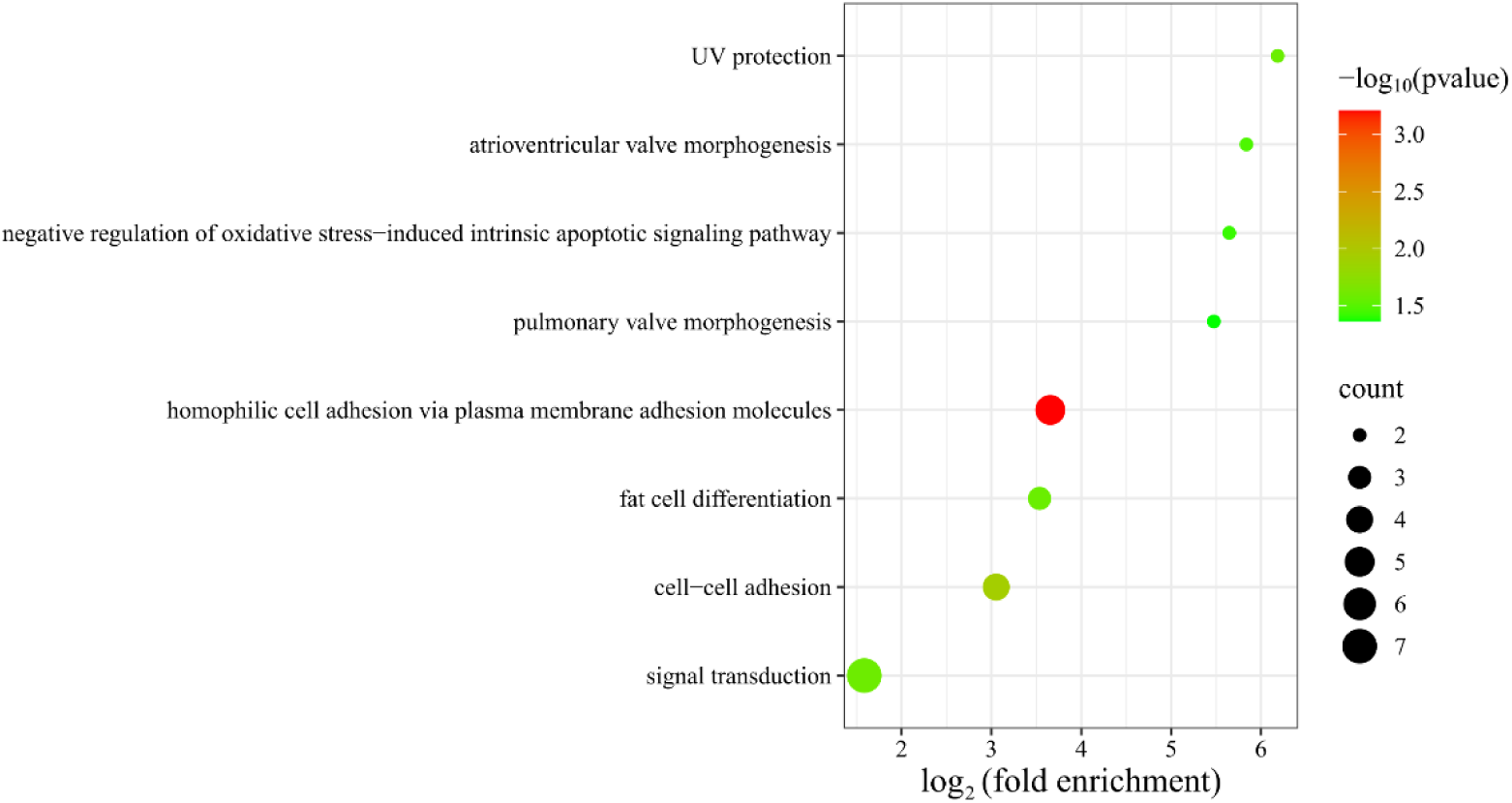
Enrichment analysis of biological processes for the identified differential proteins in the green tea group (*p* < 0.05).

#### 3.1.2 Comparison Before and After Oolong Tea Consumption

By comparing the urinary proteins of rats before and after oolong tea consumption, a total of 108 differential proteins were identified under the screening conditions of FC ≥ 1.5 or ≤ 0.67 and *p* < 0.05 by two-tailed paired *t*-test analysis. Detailed information on these differential proteins is listed in Table S1. Results from the randomized grouping test showed that the average number of differential proteins yielded was 44.60, indicating that at least 58.70% of these differential proteins were not randomly generated.

##### 3.1.2.1 Analysis of Differential Proteins Identified in the Oolong Tea Group

CCN family member 1 (FC = 0.04, *p* = 3.82×10^-02^), with the second-smallest FC value, was also identified in the before-and-after comparison of the oolong tea group.

Catalase (FC = 0, *p* = 4.60×10^-02^) exhibited a change from presence to absence. This means that it was identified in the urinary samples of rats before oolong tea consumption but not in those after consumption. Catalase possesses the functions such as antioxidant activity and catalase activity, and is involved in biological processes including cellular detoxification of hydrogen peroxide, hydrogen peroxide catabolism, response to hydrogen peroxide, cholesterol metabolism, response to fatty acids, response to oxidative stress, and triglyceride metabolism. Studies have shown that unfermented oolong tea kombucha significantly increased the mRNA levels of catalase in HEK-293 cells^27^.

Ly6/PLAUR domain-containing protein 3 (FC = 0.04, *p* = 6.54×10^-03^) exhibited the third-smallest FC value and has been reported as a biomarker and therapeutic target for acute myeloid leukemia^28^. Protocadherin-8 (FC = 16.19, *p* = 2.11×10^-02^) exhibited the largest FC value and has been reported as a prognostic biomarker for thyroid cancer^29^. High expression of *PCDH8* can also be used as a biomarker for poor prognosis in gastric cancer^30^.

##### 3.1.2.2 Analysis of Biological Pathways Enriched from Differential Proteins in the Oolong Tea Group

The differential proteins identified before and after oolong tea consumption were mainly involved in biological processes such as establishment of skin barrier, triglyceride metabolism, positive regulation of phosphatidylinositol 3-kinase/protein kinase B signal transduction, UV protection, and response to oxidative stress (Figure 3A). Oolong tea polyphenols have been shown to ameliorate circadian rhythm disorders by regulating the circadian rhythm oscillations of both intestinal flora and the transcription of circadian clock genes. The PI3K/Akt signaling pathway was one of the Kyoto Encyclopedia of Genes and Genomes (KEGG) pathways that enriched the most differentially expressed genes after oolong tea polyphenol intervention^31^. KEGG enrichment analysis revealed significant enrichment in pathways including carbon metabolism, folate biosynthesis, 2-Oxocarboxylic acid metabolism, tryptophan metabolism, and immune response (Figure 3B). Studies have shown that high consumption of oolong tea during pregnancy is associated with lower serum folate levels^32^.

**Figure 3.**
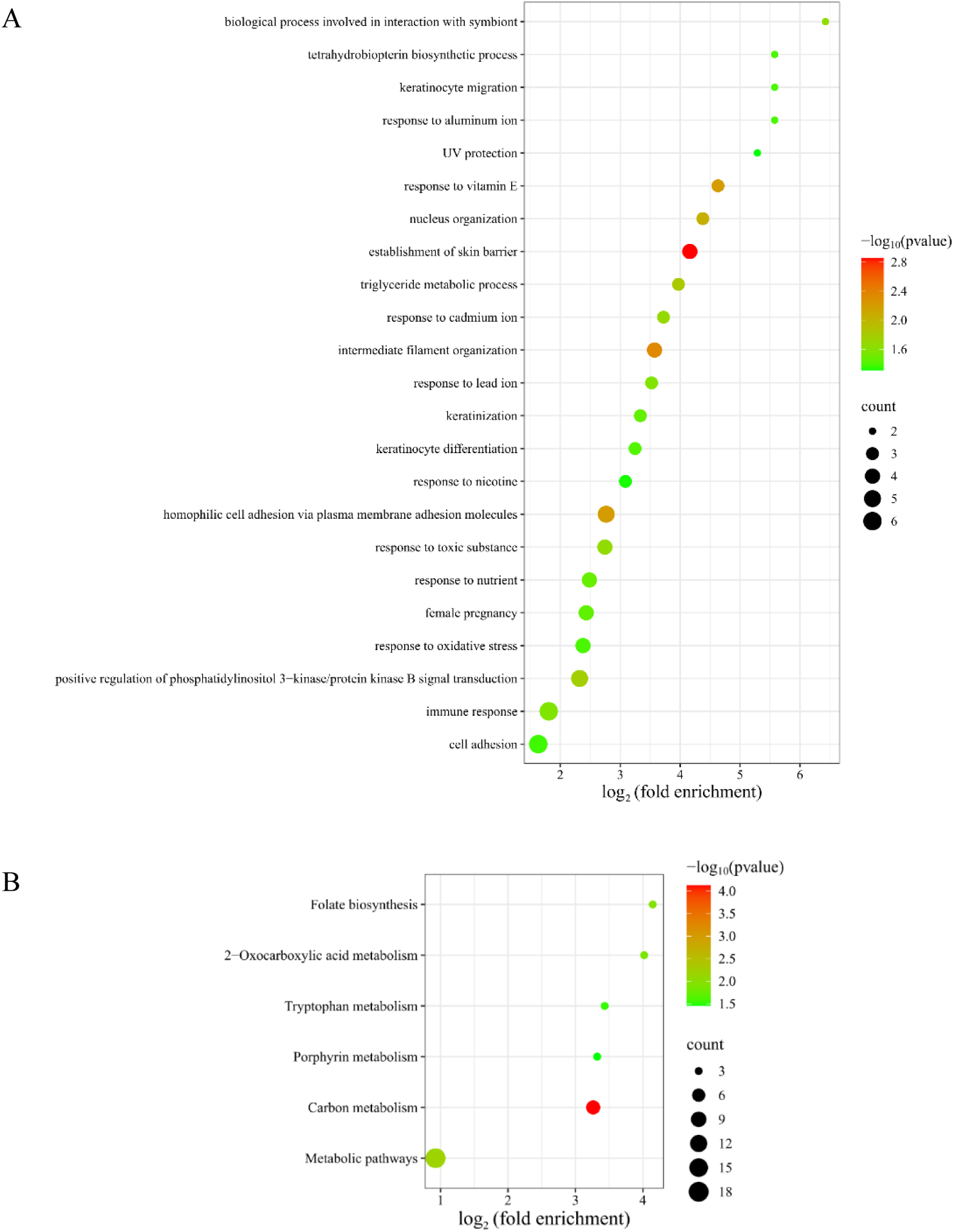
Functional annotation of identified differential proteins in the oolong tea group (*p* < 0.05): (A) biological processes; (B) KEGG pathways.

#### 3.1.3 Comparison Before and After Black Tea Consumption

By comparing the urinary proteins of rats before and after black tea consumption, a total of 142 differential proteins were identified under the screening conditions of FC ≥ 1.5 or ≤ 0.67 and *p* < 0.05 by two-tailed paired *t*-test analysis. Detailed information on these differential proteins is listed in Table S1. Results from the randomized grouping test showed that the average number of differential proteins yielded was 40.44, indicating that at least 71.52% of these differential proteins were not randomly generated.

##### 3.1.3.1 Analysis of Differential Proteins Identified in the Black Tea Group

Among the differential proteins identified before and after black tea consumption, cytoplasmic FMR1-interacting protein exhibited a change from absence to presence. This means that it was identified in the urinary samples of rats after black tea consumption but not in those before consumption. Cytoplasmic FMR1-interacting protein has been reported as a potential biomarker for diagnosing nasopharyngeal carcinoma, monitoring disease progression, and guiding therapeutic regimen selection^33^, as well as a diagnostic and prognostic biomarker for acute lymphoblastic leukemia^34^.

Acid phosphatase (FC = 29.17, *p* = 1.51×10^-02^) exhibited the second-largest FC value and is involved in biological processes such as adenosine metabolism, nucleotide metabolism, regulation of sensory perception of pain, and thiamine metabolism.

Spectrin beta chain (FC = 0.03, *p* = 1.56×10^-02^) exhibited the smallest FC value and participates in biological processes including central nervous system development, central nervous system formation, and positive regulation of interleukin-2 production. It has been reported as a biomarker for early diagnosis and precision treatment of renal clear cell carcinoma^35^.

Fibulin-1 (FC = 0.53, *p* = 8.93×10^-05^) exhibited the smallest *p*-value. Studies have shown that Fibulin-1 is closely associated with the extent of target organ damage in patients at high risk of cardiovascular disease and may serve as a biomarker for risk stratification in these patients^36^. Plasma FBLN1 levels can also be used as a biomarker to predict thyroid-related eye disease activity^37^.

##### 3.1.3.2 Analysis of Biological Pathways Enriched from Differential Proteins in the Black Tea Group

The differential proteins identified before and after black tea consumption were mainly involved in biological processes such as glyceraldehyde-3-phosphate metabolism, axon guidance, glyceraldehyde-3-phosphate biosynthesis, response to lipopolysaccharide, glycerol catabolism, response to oxidative stress, gluconeogenesis, canonical glycolysis, lipid metabolism, glucose metabolism, and fatty acid transport (Figure 4A). KEGG enrichment analysis revealed significant enrichment in pathways including the PI3K-Akt signaling pathway, fructose and mannose metabolism, carbon metabolism, inositol phosphate metabolism, glycolysis/gluconeogenesis, and tyrosine metabolism (Figure 4B). Black tea extract has been shown to inhibit growth and induce apoptosis in HepG2 cells via the PI3K-Akt signaling pathway^38^.

**Figure 4.**
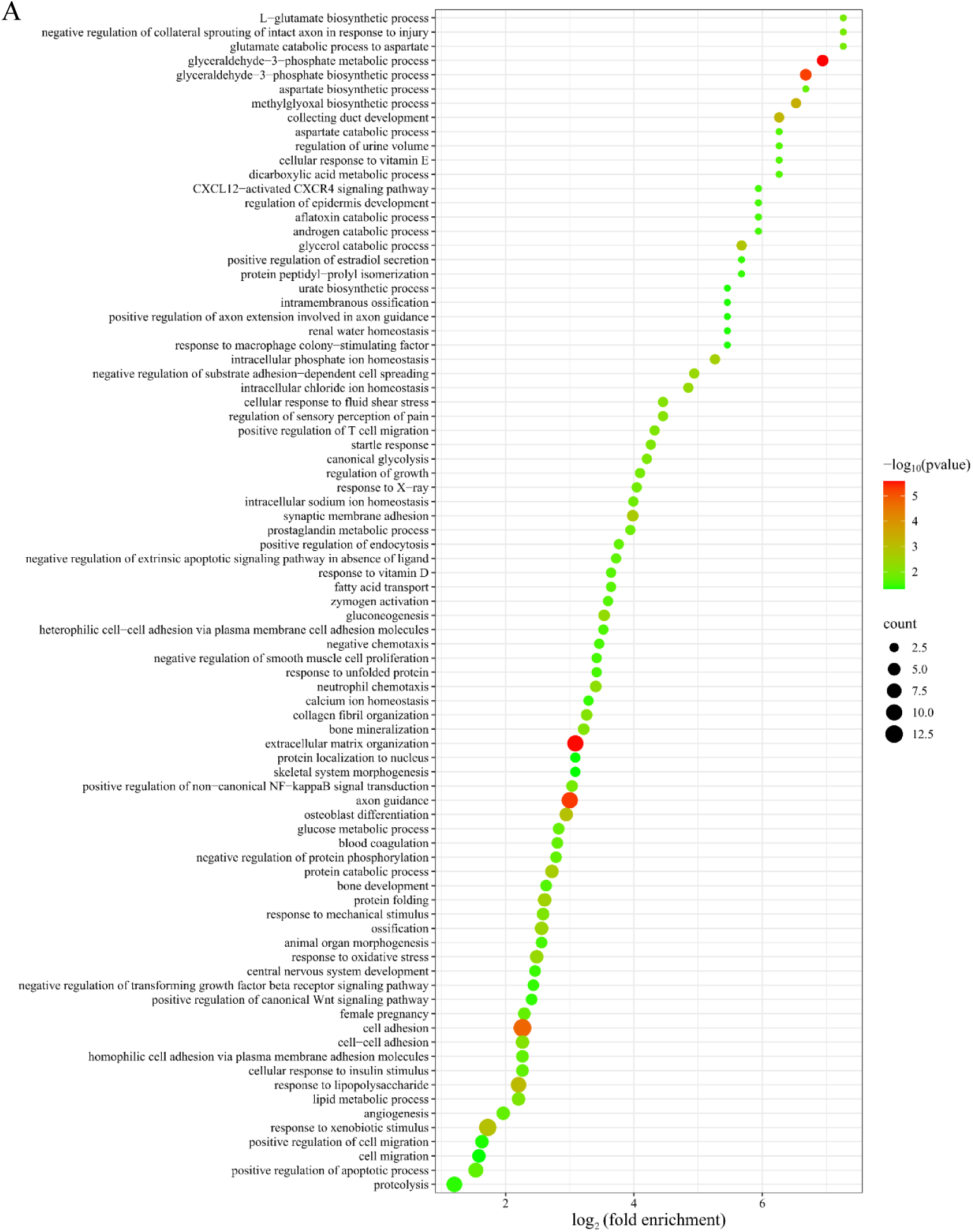
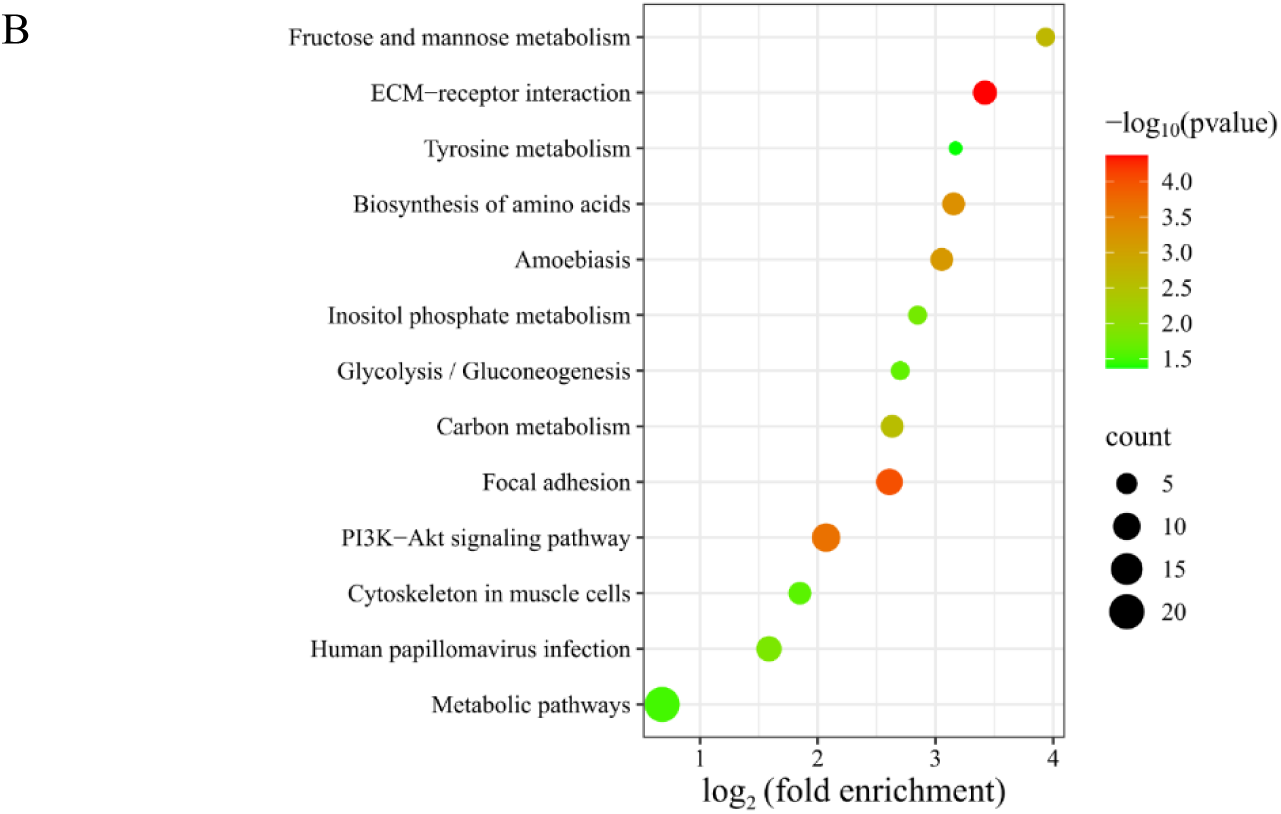
Functional annotation of identified differential proteins in the black tea group (*p* < 0.05): (A) biological processes; (B) KEGG pathways.

#### 3.1.4 Comparison Before and After Pu-erh Tea Consumption

By comparing the urinary proteins of rats before and after Pu-erh tea consumption, a total of 200 differential proteins were identified under the screening conditions of FC ≥ 1.5 or ≤ 0.67 and *p* < 0.05 by two-tailed paired *t*-test analysis. Detailed information on these differential proteins is listed in Table S1. Results from the randomized grouping test showed that the average number of differential proteins yielded was 41.91, indicating that at least 79.05% of these differential proteins were not randomly generated.

##### 3.1.4.1 Analysis of Differential Proteins Identified in the Pu-erh Tea Group

Among the differential proteins identified before and after Pu-erh tea consumption, eight proteins—including charged multivesicular body protein 2B, SPARC-like protein 1, syntaxin-7, shisa family member 7, CMRF35-like molecule 1, nebulin, proteasome activator complex subunit 1, and latent transforming growth factor beta binding protein 2—showed changes from presence to absence. This means that they were identified in the urinary samples of rats before Pu-erh tea consumption but not in those after consumption.

Charged multivesicular body protein 2B (FC = 0, *p* = 9.58 ×10^-04^) is involved in biological processes such as autophagosome maturation, autophagy, and cognition. SPARC-like protein 1 (FC = 0, *p* = 8.43 ×10^-03^) has been reported as a biomarker for human glioma progression^39^, and as a potential prognostic biomarker for fatal COVID-19 pneumonia^40^. Syntaxin-7 (FC = 0, *p* = 1.50 ×10^-02^) is involved in biological processes such as positive regulation of receptor localization to synapse and positive regulation of T cell mediated cytotoxicity. Shisa family member 7 (FC = 0, *p* = 1.78×10^-02^) possesses the functions of GABA receptor binding and ionotropic glutamate receptor binding, and is involved in gamma-aminobutyric acid receptor clustering, gamma-aminobutyric acid signaling pathway, memory, positive regulation of long-term synaptic potentiation, and regulation of AMPA glutamate receptor clustering. CMRF35-like molecule 1 (FC = 0, *p* = 2.93 ×10^-02^) is involved in immune system process. Nebulin (FC = 0, p = 2.93 × 10-02) is involved in the assembly of cardiac muscle thin filament. Proteasome activator complex subunit 1 (FC = 0, *p* = 4.02 ×10^-02^) has been reported as an independent prognostic biomarker for soft-tissue smooth muscle sarcoma^41^, and as a biomarker for rheumatoid arthritis^42^. Latent transforming growth factor beta binding protein 2 (FC = 0, *p* = 4.19×10^-02^) possesses the function of calcium ion binding and has been reported as a potential biomarker for the diagnosis and treatment of hepatocellular carcinoma^43^, a diagnostic biomarker and potential therapeutic target for pancreatic cancer^44^, and circulating LTBP-2 has also been reported as a biomarker for predicting adverse outcomes in dilated cardiomyopathy^45^.

##### 3.1.4.2 Analysis of Biological Pathways Enriched from Differential Proteins in the Pu-erh Tea Group

The differential proteins identified before and after Pu-erh tea consumption were mainly involved in biological processes such as positive regulation of phosphatidylinositol 3-kinase/protein kinase B signal transduction, axon guidance, response to hypoxia, angiogenesis, response to lipopolysaccharide, immune response, and coronary vein morphogenesis (Figure 5A).

**Figure 5.**
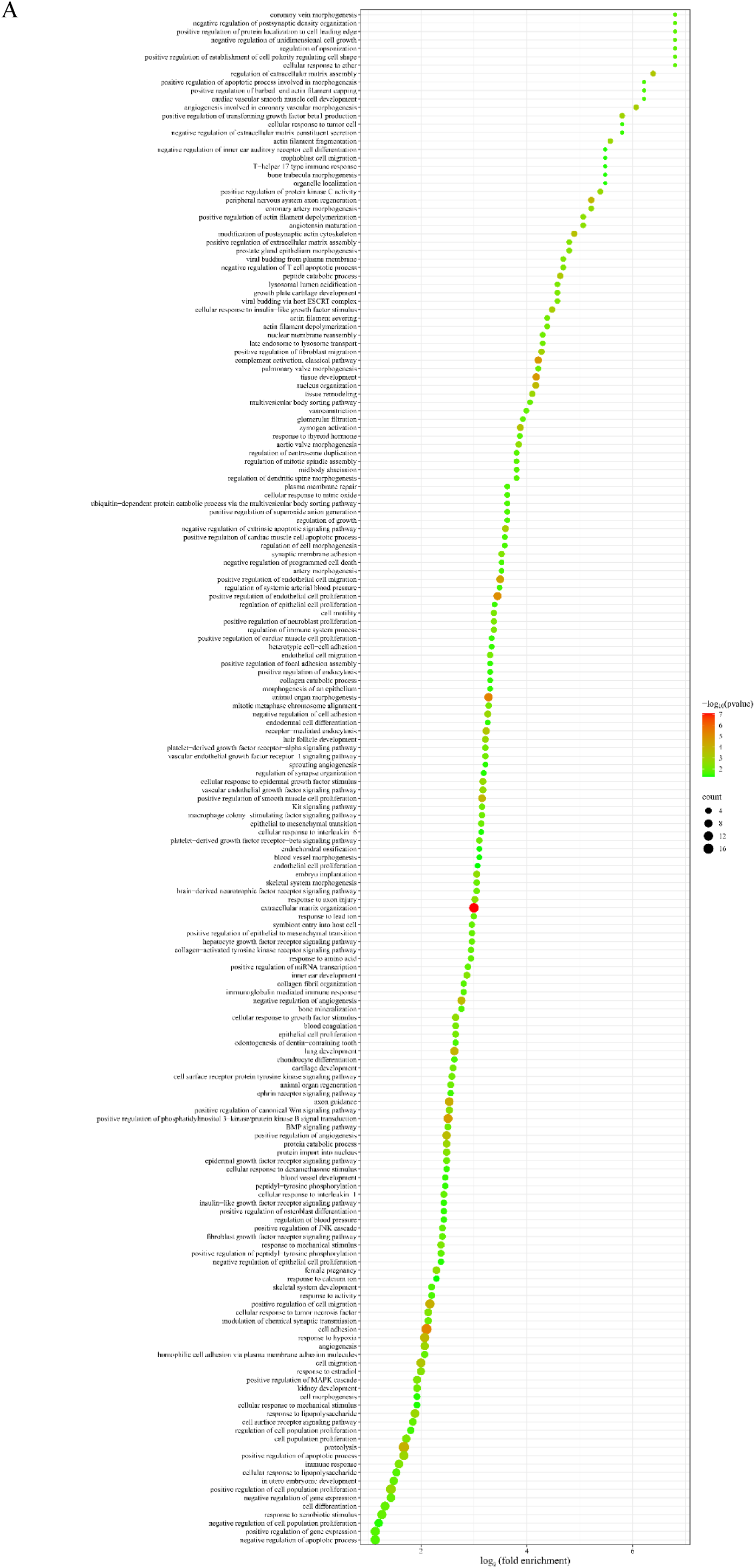
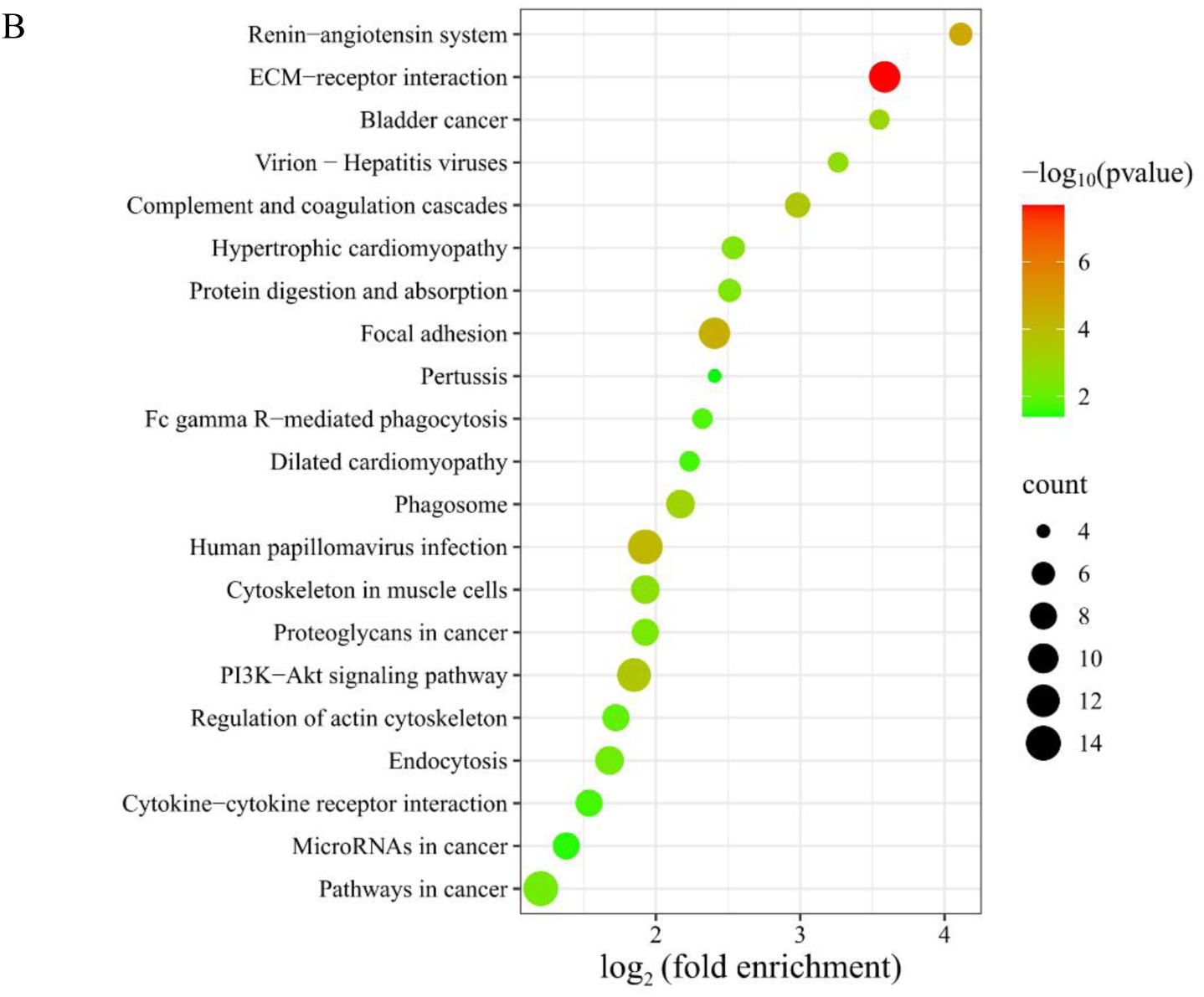
Functional annotation of identified differential proteins in the Pu-erh tea group (*p* < 0.05): (A) biological processes; (B) KEGG pathways.

KEGG enrichment analysis revealed significant enrichment in pathways including ECM-receptor interaction, renin-angiotensin system, focal adhesion, human papillomavirus infection, PI3K-Akt signaling pathway, pathways in cancer, hypertrophic cardiomyopathy, and dilated cardiomyopathy (Figure 5B). Tea fucoxanthin, a bioactive component of black tea, has been reported to regulate glycolipid metabolism through the IRS-1/PI3K/Akt signaling pathway^46^.

#### 3.1.5 Comparison Before and After Black Coffee Consumption

By comparing the urinary proteins of rats before and after black coffee consumption, a total of 246 differential proteins were identified under the screening conditions of FC ≥ 1.5 or ≤ 0.67 and *p* < 0.05 by two-tailed paired *t*-test analysis. Detailed information on these differential proteins is listed in Table S1. Results from the randomized grouping test showed that the average number of differential proteins yielded was 44.96, indicating that at least 81.72% of these differential proteins were not randomly generated.

##### 3.1.5.1 Analysis of Differential Proteins Identified in the Black Coffee Group

Among the differential proteins identified before and after black coffee consumption, ten proteins—including tetraspanin, RAB5B, shisa family member 7, latent transforming growth factor beta binding protein 2, tumor necrosis factor receptor superfamily member 4, choline transporter-like protein 2, glycoprotein hormone alpha-2, protocadherin alpha-4, CCN family member 1, and tissue-type plasminogen activator—showed changes from presence to absence. This means that they were identified in the urinary samples of rats before black coffee consumption but not in those after consumption.

Tetraspanin (FC = 0, *p* = 2.39×10^-04^) is involved in the negative regulation of blood coagulation, regulation of gene expression, and spermatogenesis. It has been reported as a diagnostic and prognostic biomarker as well as a therapeutic target in colorectal cancer^47^.

RAB5B (FC = 0, *p* = 5.11×10^-04^) possesses the functions such as GDP binding, GTP binding, and GTPase activity, and is involved in biological processes such as antigen processing and presentation, and endocytosis. Shisa family member 7 (FC = 0, *p* = 1.49×10^-03^) possesses the functions such as GABA receptor binding, ionotropic glutamate receptor binding, gamma-aminobutyric acid receptor clustering, gamma-aminobutyric acid signaling pathway, memory, positive regulation of long-term synaptic potentiation, and regulation of AMPA glutamate receptor clustering.

Latent transforming growth factor beta binding protein 2 (FC = 0, *p* = 3.95×10^-03^) has been reported as a potential biomarker for the diagnosis and treatment of hepatocellular carcinoma^43^, a diagnostic biomarker and potential therapeutic target for pancreatic cancer^44^, and circulating LTBP-2 has also been reported as a biomarker for predicting adverse outcomes in dilated cardiomyopathy^45^.

Tumor necrosis factor receptor superfamily member 4 (FC = 0, *p* = 5.66×10^-03^) is involved in biological processes such as cellular defense response, inflammatory response, negative regulation of activation-induced cell death of T cells, positive regulation of B cell proliferation, T cell proliferation, and regulation of apoptotic process. *TNFRSF4* has been reported as a potential biomarker for prognosis and immunomodulation in endometrial cancer^48^, and as an independent potential biomarker for prognostic prediction in pancreatic cancer^49^.

Choline transporter-like protein 2 (FC = 0, *p* = 1.31×10^-02^) is involved in biological processes such as choline transport and ethanolamine transport. Glycoprotein hormone alpha-2 (FC = 0, *p* = 3.49×10^-02^) participates in the adenylate cyclase-activating G protein-coupled receptor signaling pathway and cell surface receptor signaling pathway. Tissue-type plasminogen activator (FC = 0, *p* = 4.53×10^-02^) is involved in biological processes including the regulation of synaptic plasticity, glutamatergic synaptic transmission, trans-synaptic signaling by BDNF, and responses to cAMP, fatty acids, and hypoxia. CCN family member 1 (FC = 0, *p* = 4.43×10^-02^) has been reported as an early biomarker of infarct size and left ventricular dysfunction in patients with ST-segment elevation myocardial infarction^22^.

Matrilin 2 (FC = 0.26, *p* = 3.24×10^-05^) exhibited the smallest *p*-value and features a calcium-binding function. It is involved in axon guidance, dendrite regeneration, glial cell migration, neuron migration, neuron projection development, and response to axon injury. Matrilin 2 has been reported as a specific biomarker to differentiate inert from clinically aggressive pilocytic astrocytoma^50^ and serves as a prognostic biomarker for osteosarcoma^51^.

##### 3.1.5.2 Analysis of Biological Pathways Enriched from Differential Proteins in the Black Coffee Group

The differential proteins identified before and after black coffee consumption were mainly involved in biological processes such as axon guidance, positive regulation of phosphatidylinositol 3-kinase/protein kinase B signal transduction, response to lipopolysaccharide, ephrin receptor signaling pathway, complement activation, heart development, negative regulation of apoptotic process, and immune response (Figure 6A). KEGG enrichment analysis revealed significant enrichment in pathways including complement and coagulation cascades, focal adhesion, PI3K-Akt signaling pathway, human papillomavirus infection, nitrogen metabolism, platelet activation, microRNAs in cancer, fluid shear stress and atherosclerosis, and pathways in cancer (Figure 6B). Coffee consumption has been reported as one of the risk factors for human papillomavirus (HPV) infection^52^. Caffeine-mediated inhibition of the DNA damage response (DDR) reduces viral genome replication during the early stages of HPV infection, whereas DDR inhibition may lead to an increase in viral amplicon replication during the maintenance phase^53^. In addition, caffeine activates the PI3K/Akt pathway and inhibits apoptosis in the SH-SY5Y cell model of Parkinson’s disease^54^.

**Figure 6.**
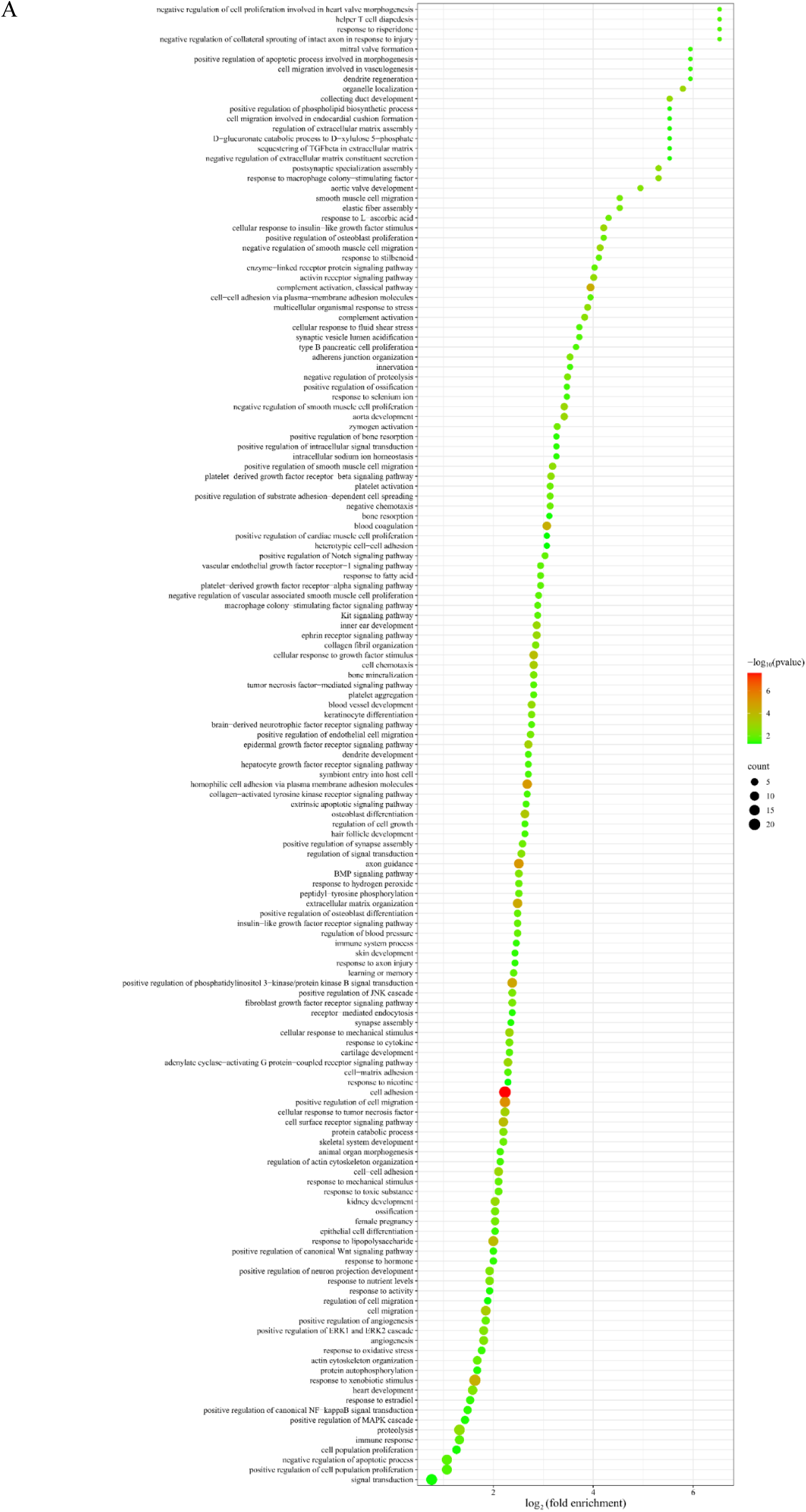
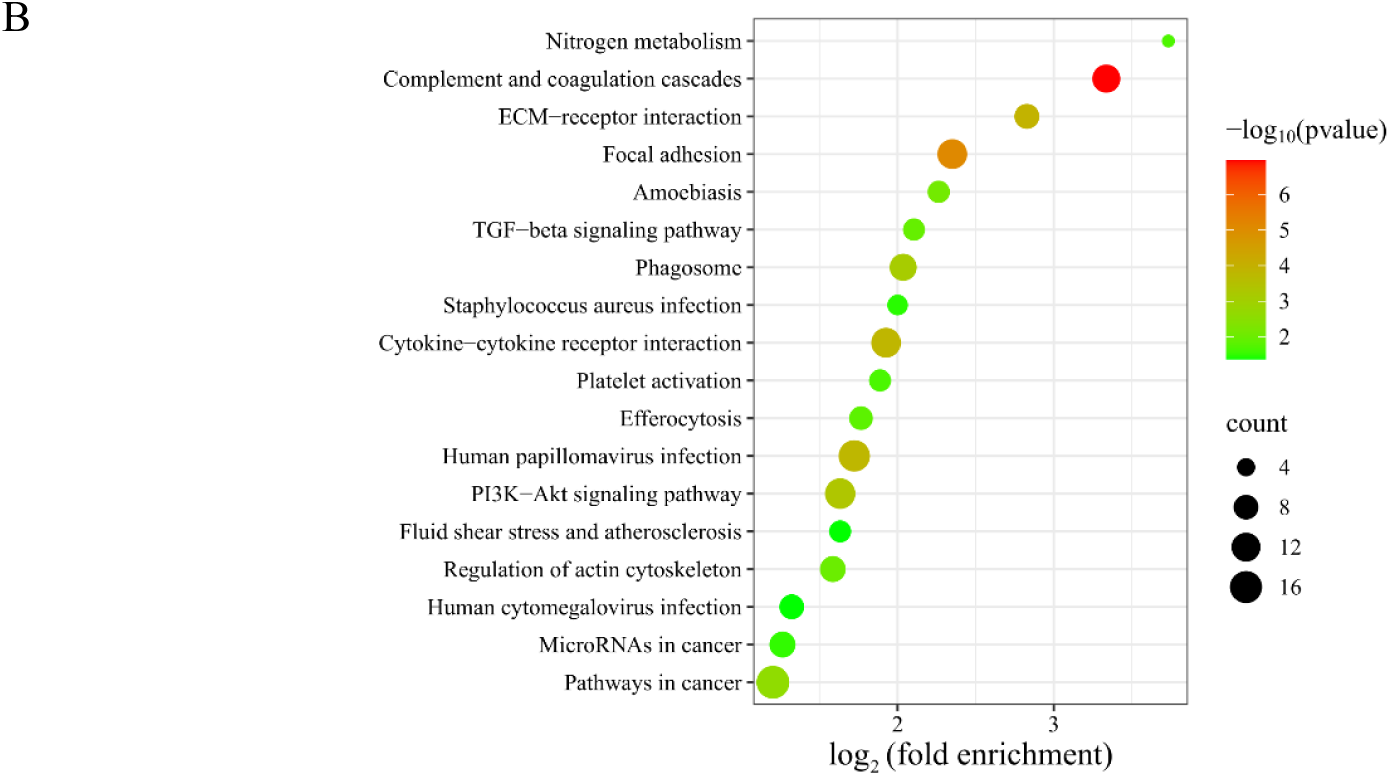
Functional annotation of identified differential proteins in the black coffee group (*p* < 0.05): (A) biological processes; (B) KEGG pathways.

#### 3.1.6 Before-and-After Comparison in the Control Group

By comparing the urinary proteins of rats before and after sterile water consumption, a total of 83 differential proteins were identified under the screening conditions of FC ≥ 1.5 or ≤ 0.67 and *p* < 0.05 by two-tailed paired *t*-test analysis. Detailed information on these differential proteins is listed in Table S1. Results from the randomized grouping test showed that the average number of differential proteins yielded was 40.90, indicating that at least 50.72% of these differential proteins were not randomly generated. These differential proteins were mainly involved in biological processes such as skeletal system morphogenesis, heart development, and cartilage development. Although the before-and-after comparisons minimized the effects of individual differences, they could not avoid the effects of growth and development. Therefore, to exclude the effects of short-term growth and development, between-group comparisons of urinary proteins were performed in rats from the experimental and control groups after consuming teas, black coffee, or sterile water.

### 3.2 Comparative Analysis Between Groups After Consuming Teas and Black Coffee

#### 3.2.1 Comparison Between the Green Tea Group and the Control Group

By comparing the urinary proteins of rats in the green tea group with those in the control group, a total of 59 differential proteins were identified under the screening conditions of FC ≥ 1.5 or ≤ 0.67 and *p* < 0.05 by two-tailed unpaired *t*-test analysis. Detailed information on these differential proteins is listed in Table S2. Results from the randomized grouping test showed that the average number of differential proteins yielded was 37.69, indicating that at least 36.12% of these differential proteins were not randomly generated.

##### 3.2.1.1 Analysis of Differential Proteins Identified by Comparison between the Green Tea Group and the Control Group

Suppressor of tumorigenicity 14 protein homolog (FC = ∞, *p* = 2.78×10^-02^) exhibited a change from absence to presence. This means that it was identified in the green tea group samples but not in any of the control group samples. It has been reported to play an important role in all stages of epithelial tumorigenesis and development, and holds potential as a target for anticancer therapy, as well as a diagnostic and prognostic marker^55, 56^.

Myosin light chain 12A (FC = 152.50, p = 4.76 × 10⁻²) exhibited the second-largest FC value among the identified differential proteins. It possesses the functions such as calcium ion binding. *MYL12A* is highly expressed in smooth and skeletal muscle tissues and encodes a non-sarcomeric myosin regulatory light chain protein that regulates muscle contraction. It is considered a new candidate gene related to muscle strength^57^. In addition, Myl9/12 has been reported to be involved in the pathogenesis of inflammatory bowel disease and may serve as a new therapeutic target. Plasma Myl9 levels may also serve as a biomarker for inflammatory bowel disease^58^. Another study showed that cardiac-specific overexpression of MYL12A rescued diminished cardiac contractility in necdin-deficient mice^59^.

Metalloproteinase inhibitor 3 (TIMP-3) (FC = 25.42, *p* = 3.66×10^-02^) exhibited the third-largest FC value. It possesses the functions such as metalloendopeptidase inhibitor activity and zinc ion binding. It is involved in biological processes including mesenchymal cell differentiation in bone development, cellular response to hypoxia and interleukin-6, and negative regulation of vascular associated smooth muscle cell proliferation. TIMP-3 can be used as a surrogate marker for the response to green tea polyphenols (GTPs) and their major component epigallocatechin-3-gallate (EGCG). GTP and EGCG inhibit prostate cancer cell migration and invasion by reactivating TIMP-3 and inhibiting MMP-2/MMP-9 activity^60^. They also induce TIMP-3 expression in breast cancer cells, delaying progression and invasion^61^. Green tea extract restores circadian expression of *TIMP3* disrupted by UVB irradiation^62^. Furthermore, TIMP-3 has been reported as a cancer biomarker and potential therapeutic target^63^. For example, plasma TIMP3 levels in patients with oral squamous cell carcinoma were significantly lower than in healthy controls, and plasma TIMP3 could be used as a potential biomarker for predicting the tumor stage and T status in patients with oral squamous cell carcinoma^64^.

Crk-like protein (FC = 23.87, *p* = 4.4×10^-02^) exhibited the fourth-largest FC value and is involved in biological processes such as blood vessel development, B cell apoptosis, and lipid metabolism. Studies have shown that CrkL can be proposed as a soluble serum biomarker in patients with breast cancer, especially in the advanced disease stages^65^.

Phospholipase B1 (FC = 0.14, *p* = 2.51×10^-02^) exhibited the smallest FC value and participates in triglyceride catabolic process, retinol metabolic process, and phospholipid metabolic process. It has been identified as a potential antigen of glioblastoma, correlating with patient survival and infiltration of antigen-presenting cells, and is a potential target for glioblastoma mRNA vaccine development^66^. Another study showed that the rs117512489 variant of *PLB1* was significantly associated with survival in patients with non-small cell lung cancer^67^. *PLB1* has also been reported as a candidate risk gene for rheumatoid arthritis^68^ and possibly for juvenile dermatomyositis^69^.

Lithostathine (FC = 0.40, *p* = 2.52×10^-02^) exhibited the third-smallest FC value and possesses the functions such as growth factor activity, phosphatase binding, signaling receptor activity, and signaling receptor binding. It is involved in biological processes such as antimicrobial humoral immune response mediated by antimicrobial peptide, cellular response to chemokine, cellular response to gastrin, liver regeneration, and response to hypoxia. Lithostathine has been reported to be involved in the pathophysiologic processes of Alzheimer’s disease, with a significant increase in the early stages of the disease and persistently high levels throughout the course of the disease^70^.

##### 3.2.1.2 Analysis of Biological Pathways Enriched from Differential Proteins

Identified by Comparison between the Green Tea Group and the Control Group The differential proteins identified by comparison between the green tea group and the control group were mainly involved in biological processes such as axon guidance, ephrin receptor signaling pathway, cell migration, positive regulation of ERK1 and ERK2 cascade, angiogenesis, aorta development, semaphorin-plexin signaling pathway, neural crest cell migration, heart development, positive regulation of phosphatidylinositol 3-kinase/protein kinase B signal transduction, and sphingosine biosynthesis (Figure 7A). EGCG, the major catechin in green tea, has been reported to inhibit angiogenesis through a variety of mechanisms, including induction of apoptosis and promotion of cell cycle arrest, modulation of miRNA expression profile, and inhibition of vascular endothelial growth factor (VEGF) binding to its receptor^71^. Sphingosine concentrations in plasma have been found to significantly increased 2 hours after ingestion of green tea extract^72^.

**Figure 7.**
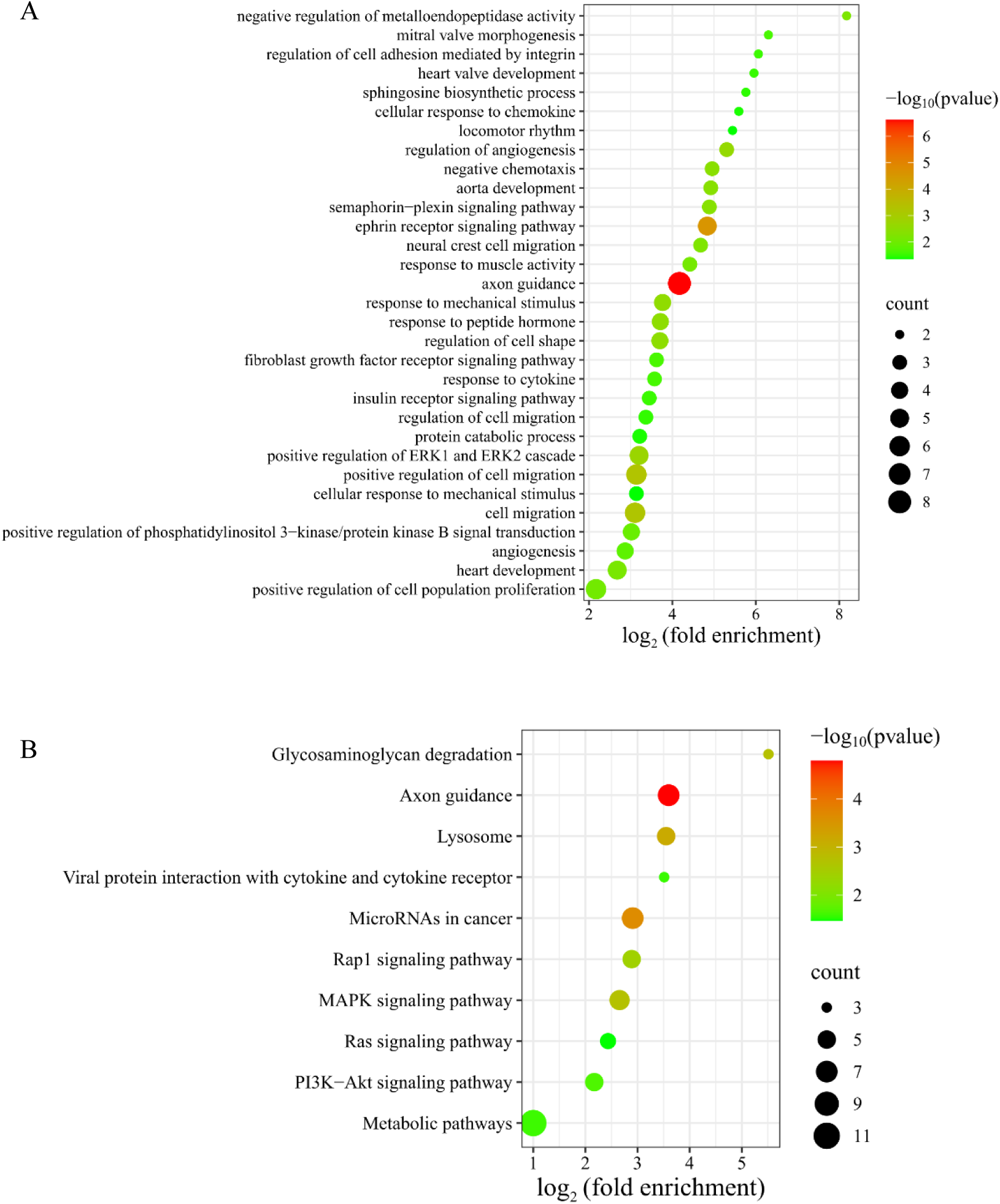
Functional annotation of differential proteins identified in the comparison between the green tea group and the control group (*p* < 0.05): (A) biological processes; (B) KEGG pathways.

KEGG enrichment analysis revealed significant enrichment in pathways including axon guidance, microRNAs in cancer, lysosome, MAPK signaling pathway, PI3K-Akt signaling pathway, and metabolic pathways (Figure 7B). Proanthocyanidins, important polyphenols in tea, have been shown to regulate miRNAs involved in cancer, glucose and lipid homeostasis^73^. Tea polysaccharides, bioactive components of green tea, target lysosomes and induce apoptosis via a lysosomal-mitochondrial pathway mediated caspase cascade, inhibiting proliferation of colon cancer cell line CT26^74^. EGCG, an active ingredient in green tea, promotes the expression of protein kinase C alpha (PRKCA) and attenuates lipopolysaccharide-induced acute lung injury and inflammatory responses. This may be related to PRKCA regulating the MAPK signaling pathway and influencing the release of pro-inflammatory cytokines from macrophages^75^. Green tea extract inhibits HepG2 cell growth and induces apoptosis via the PI3K/Akt pathway^38^. Additionally, EGCG inhibits cardiomyocyte apoptosis, restores autophagic flux, and attenuates myocardial ischemia/reperfusion injury through the PI3K/Akt signaling pathway^76^.

#### 3.2.2 Comparison Between the Oolong Tea Group and the Control Group

By comparing the urinary proteins of rats in the oolong tea group with those in the control group, a total of 60 differential proteins were identified under the screening conditions of FC ≥ 1.5 or ≤ 0.67 and *p* < 0.05 by two-tailed unpaired *t*-test analysis. Detailed information on these differential proteins is listed in Table S2. Results from the randomized grouping test showed that the average number of differential proteins yielded was 40.48, indicating that at least 32.53% of these differential proteins were not randomly generated.

##### 3.2.2.1 Analysis of Differential Proteins Identified by Comparison between the Oolong Tea Group and the Control Group

Selenoprotein F (FC = ∞, *p* = 2.99×10^-02^) exhibited a change from absence to presence. This means that it was identified in the oolong tea group samples but not in any of the control group samples. *SELENOF* gene polymorphisms and SELENOF dysregulation are closely related to diseases including cancer and neurodegenerative diseases. Since SELENOF is sensitive to selenium, it may serve as a therapeutic target in the pathological processes of related diseases^77^.

Endoribonuclease LACTB2 (FC = 3690.17, *p* = 4.69×10^-02^) exhibited the second-largest FC value among the identified differential proteins. LACTB2 has been reported as a biomarker for predicting the prognosis of radiotherapy in nasopharyngeal carcinoma and as a potential therapeutic target for improving the radiosensitivity of nasopharyngeal carcinoma^78^.

Myosin light chain 12A and phospholipase B1 were also identified as differential proteins by comparison between the oolong tea group and the control group. Myosin light chain 12A (FC = 487.56, *p* = 1.85×10^-02^) exhibited the third-largest FC value, while phospholipase B1 (FC = 0.25, *p* = 3.75×10^-02^) exhibited the fifth-smallest FC value.

Ectonucleotide pyrophosphatase/phosphodiesterase 2 (FC = 12.84, *p* = 1.58×10^-02^) exhibited the fifth-largest FC value. ENPP2 has been reported as a potential biomarker for breast cancer initiation and progression^79^, as well as for the diagnosis and prognosis of hepatocellular carcinoma^80^. It is also considered an independent prognostic marker for predicting biochemical recurrence in prostate cancer patients with postoperative prostate specific antigen levels ≥ 0.2 ng/mL^81^.

Type I keratin KA11 (FC = 0.04, *p* = 3.54×10^-02^) exhibited the smallest FC value and is involved in biological processes such as inflammatory response, establishment of skin barrier, keratinization, and morphogenesis of an epithelium. Dmx-like 1 (FC = 16.21, *p* = 2.41×10^-02^) exhibited the fourth-largest FC value and is involved in vacuolar acidification.

Von Willebrand factor (FC = 4.94, *p* = 6.29×10^-04^) exhibited the smallest *p*-value. It has been identified as a biomarker for diagnosing clinically significant portal hypertension (CSPH) and severe portal hypertension (SPH) in patients with cirrhotic, showing a moderate correlation with hepatic venous pressure gradient (HVPG) measurements^82^. VWF is also a biomarker for assessing the severity of valvular heart disease, evaluating treatment efficacy, and predicting patient prognosis^83^. Furthermore, VWF also serves as a biomarker for predicting the risk of venous thromboembolism, and is a potential target for its prevention and treatment^84^. Additionally, VWF has been reported as a novel prognostic biomarker for lung adenocarcinoma^85^.

##### 3.2.2.2 Analysis of Biological Pathways Enriched from Differential Proteins Identified by Comparison between the Oolong Tea Group and the Control Group

The differential proteins identified by comparison between the oolong tea group and the control group were mainly involved in biological processes such as intermediate filament organization, keratinization, peptide cross-linking, animal organ regeneration, complement activation, morphogenesis of an epithelium, response to lipopolysaccharide, positive regulation of coagulation, and phosphatidylcholine catabolism (Figure 8A). Studies have shown that administration of oolong tea polyphenols in mice reduces serum lipopolysaccharide levels, alleviates lipopolysaccharide-induced microglial activation, improves neuroinflammation and neuronal damage, and reduces elevated levels of the neurotoxic metabolite glutamate^86^. KEGG enrichment analysis revealed significant enrichment in pathways including lysosome, complement and coagulation cascades, PI3K-Akt signaling pathway, and *staphylococcus aureus* infection (Figure 8B). Studies have shown that the PI3K-Akt signaling pathway is one of the KEGG pathways that enriched the most differentially expressed genes following intervention with oolong tea polyphenols^31^. Additionally, water-soluble extracts from oolong tea have been shown to protect *Caenorhabditis elegans* from *Staphylococcus aureus* infection^87^.

**Figure 8.**
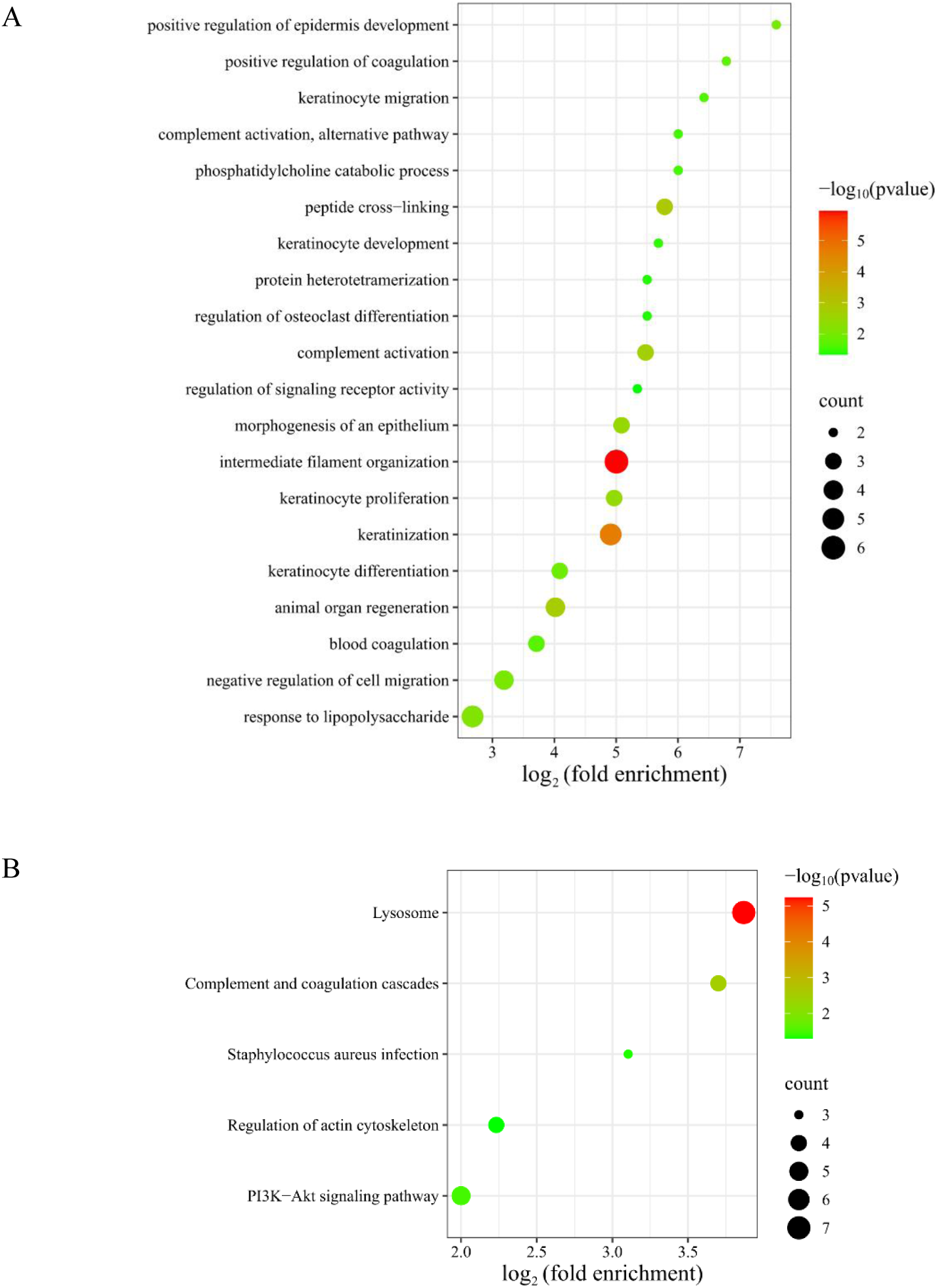
Functional annotation of differential proteins identified in the comparison between the oolong tea group and the control group (*p* < 0.05): (A) biological processes; (B) KEGG pathways.

#### 3.2.3 Comparison Between the Black Tea Group and the Control Group

By comparing the urinary proteins of rats in the black tea group with those in the control group, a total of 94 differential proteins were identified under the screening conditions of FC ≥ 1.5 or ≤ 0.67 and *p* < 0.05 by two-tailed unpaired *t*-test analysis. Detailed information on these differential proteins is listed in Table S2. Results from the randomized grouping test showed that the average number of differential proteins yielded was 41.39, indicating that at least 55.97% of these differential proteins were not randomly generated.

##### 3.2.3.1 Analysis of Differential Proteins Identified by Comparison between the Black Tea Group and the Control Group

Selenoprotein F, suppressor of tumorigenicity 14 protein homolog, Dmx-like 1 and von Willebrand factor were also identified as differential proteins by comparison between the black tea group and the control group. Selenoprotein F (FC = ∞, *p* = 9.09×10^-03^) and suppressor of tumorigenicity 14 protein homolog (FC = ∞, *p* = 1.87×10^-02^) exhibited changes from absence to presence. This means that they were identified in the black tea group samples but not in any of the control group samples. Dmx-like 1 (FC = 12.49, *p* = 2.30×10^-02^) exhibited the fifth-largest FC value among the identified differential proteins. Von Willebrand factor (FC = 4.74, *p* = 8.26×10^-04^) exhibited the fifth-smallest *p*-value.

Synaptobrevin homolog YKT6 (FC = 35815.21, *p* = 4.37×10^-02^) exhibited the third-largest FC value. Studies have shown that *YKT6* can serve as an independent prognostic biomarker and a potential immunotherapy target for oral squamous cell carcinoma^88^. Its upregulation is closely associated with the progression of hepatocellular carcinoma, and serves as a potential biomarker for poor prognosis in hepatocellular carcinoma^89^. YKT6 has also been reported as a potential prognostic and immunotherapy biomarker for cell carcinoma and endocervical adenocarcinoma^90^, as well as a potential prognostic and diagnostic biomarker for lung adenocarcinoma^91^.

PKHD1 ciliary IPT domain-containing fibrocystin/polyductin (FC = 149.77, *p* = 2.47×10^-02^) exhibited the fourth-largest FC value and has been reported as a potential prognostic biomarker for colorectal cancer^92^.

Keratin 2 (FC = 0.24, *p* = 2.33×10^-02^) exhibited the smallest FC value, and is involved in biological processes such as keratinocyte activation, development, migration, proliferation, intermediate filament organization, and keratinization.

Ephrin-A1 (FC = 4.30, *p* = 5.43×10^-05^) exhibited the smallest *p*-value. It is involved in biological processes such as angiogenesis, negative/positive regulation of MAPK cascade, aortic valve morphogenesis, and axon guidance. EFNA1 has been reported as a prognostic biomarker and potential therapeutic target for cervical cancer^93^. Serum Ephrin-A1 may serve as a diagnostic biomarker for colorectal cancer^94^.

Complement factor B (FC = 0.56, *p* = 4.32×10^-04^) exhibited the second-smallest *p*-value. It is involved in biological processes such as complement activation, positive regulation of apoptotic cell clearance, and responses to bacterium, lipopolysaccharide, and nutrient. It has been reported as a serum biomarker for pancreatic cancer^95^ and a potential biomarker and therapeutic target for lung adenocarcinoma^96^.

Interleukin 6 cytokine family signal transducer (FC = 4.47, *p* = 5.50×10^-04^) exhibited the fourth-smallest *p*-value and has been reported as a predictive and prognostic biomarker in breast cancer^97^.

Complement C1r subcomponent-like (FC = 5.32, *p* = 8.53 × 10^-03^) has been reported as a prognostic biomarker for hepatocellular carcinoma^98^ and as an independent adverse prognostic biomarker in patients with glioma, with potential as a clinical immunotherapy target^99^.

##### 3.2.3.2 Analysis of Biological Pathways Enriched from Differential Proteins Identified by Comparison between the Black Tea Group and the Control Group

The differential proteins identified by comparison between the black tea group and the control group were mainly involved in biological processes such as blood coagulation, complement activation, proteolysis, interleukin-11-mediated signaling pathway, axon guidance, and ephrin receptor signaling pathway (Figure 9A).

**Figure 9.**
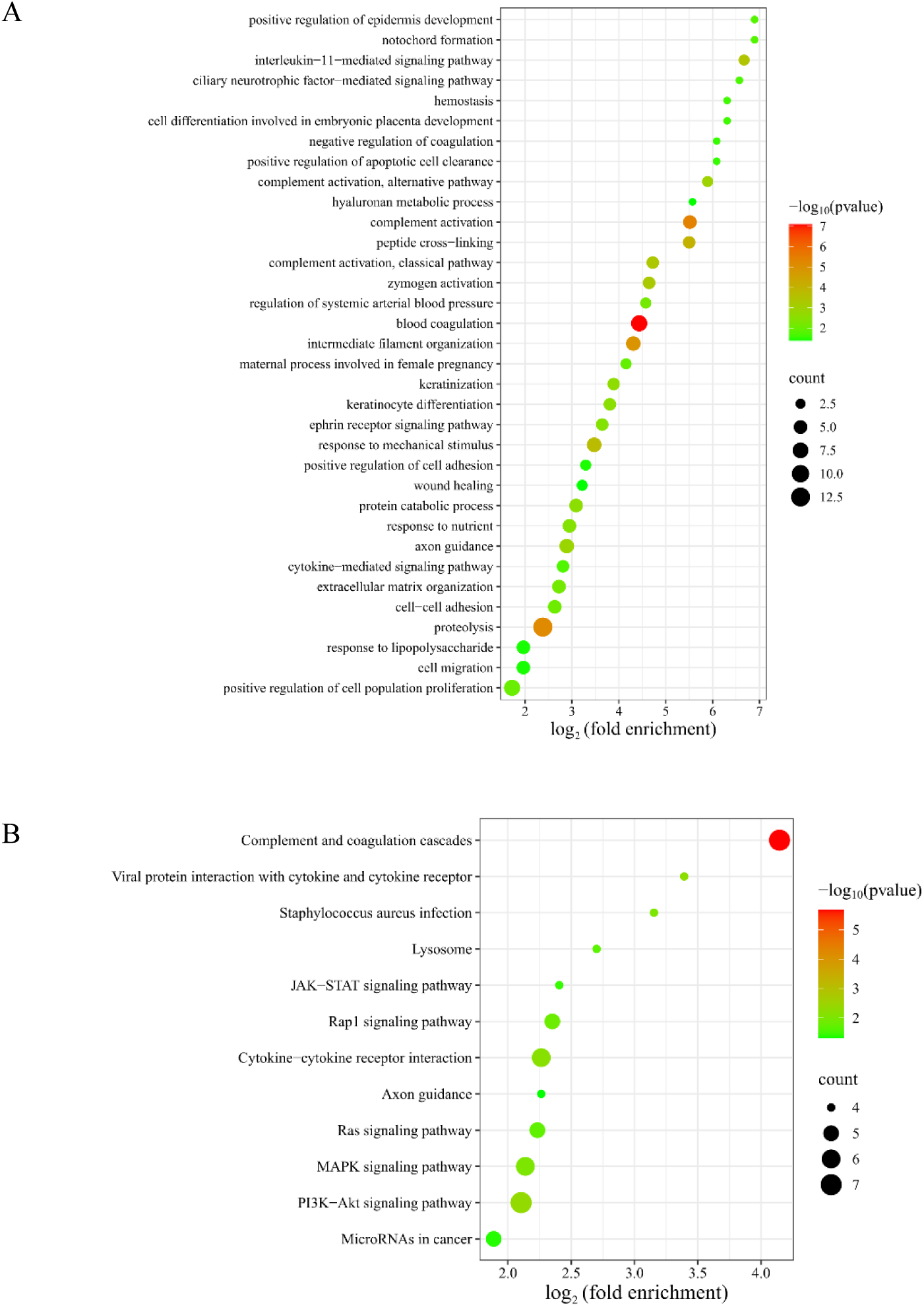
Functional annotation of differential proteins identified in the comparison between the black tea group and the control group (*p* < 0.05): (A) biological processes; (B) KEGG pathways.

KEGG enrichment analysis revealed significant enrichment in pathways including complement and coagulation cascades, PI3K-Akt signaling pathway, viral protein interaction with cytokine and cytokine receptor, MAPK signaling pathway, Ras signaling pathway, JAK-STAT signaling pathway, and microRNAs in cancer (Figure 9B). Studies have shown that black tea extract inhibits the proliferation of HepG2 cells and induces apoptosis via the PI3K-Akt signaling pathway^38^. The main polyphenols in black tea, theaflavins and thearubigins, induce apoptosis in human malignant melanoma cells (A375) through the JNK and p38 MAPK signaling pathways^100^. Additionally, theaflavins disrupt lipid rafts, preventing RET from anchoring to the cell membrane, thereby inhibiting its downstream PI3K/Akt/Bad and Ras/Raf/ERK pathways, activating the p38 MAPK/caspase-8 pathway, and inducing apoptosis in medullary thyroid cancer cells^101^. Black tea can also inhibit tumor-induced thymic involution through mechanisms such as preventing the downregulation of IL-7Rα in thymocytes, maintaining phosphorylation of JAK3 and STAT5, and protecting the JAK-STAT signaling pathway^102^.

#### 3.2.4 Comparison Between the Pu-erh Tea Group and the Control Group

By comparing the urinary proteins of rats in the Pu-erh tea group with those in the control group, a total of 95 differential proteins were identified under the screening conditions of FC ≥ 1.5 or ≤ 0.67 and *p* < 0.05 by two-tailed unpaired *t*-test analysis. Detailed information on these differential proteins is listed in Table S2. Results from the randomized grouping test showed that the average number of differential proteins yielded was 34.63, indicating that at least 63.55% of these differential proteins were not randomly generated.

##### 3.2.4.1 Analysis of Differential Proteins Identified by Comparison between the Pu-erh Tea Group and the Control Group

Suppressor of tumorigenicity 14 protein homolog, 45 kDa calcium-binding protein, myosin light chain 12A, Crk-like protein, von Willebrand factor, ephrin-A1, and complement factor B were also identified as differential proteins by comparison between the Pu-erh tea group and the control group. Suppressor of tumorigenicity 14 protein homolog (FC = ∞, *p* = 3.07×10^-02^) exhibited a change from absence to presence. This means that it was identified in the Pu-erh tea group samples but not in any of the control group samples. 45 kDa calcium-binding protein (FC = 0.07, *p* = 1.13×10^-03^) had the smallest FC value among the identified differential proteins. Myosin light chain 12A (FC = 190.12, *p* = 1.16×10^-02^) exhibited the second-largest FC value. Crk-like protein (FC = 26.23, *p* = 4.70×10^-03^) exhibited the third-largest FC value. Von Willebrand factor (FC = 3.99, *p* = 2.13×10^-02^) exhibited the fourth-largest FC value. Ephrin-A1 (FC = 3.62, *p* = 3.39×10^-04^) had the sixth-largest FC value and the fourth-smallest *p*-value. Complement factor B (FC = 0.47, *p* = 1.10×10^-05^) exhibited the second-smallest *p*-value.

Cadherin, EGF LAG seven-pass G-type receptor 2 (FC = 0.16, *p* = 3.07×10^-02^) exhibited the second-smallest FC value, possesses the functions of calcium ion binding and G protein-coupled receptor activity. It is involved in biological processes such as cerebrospinal fluid secretion, motor neuron migration, neural plate anterior/posterior regionalization, ventricular system development, and Wnt signaling pathway. It has been reported as a prognostic biomarker for hepatocellular carcinoma^103^.

Cytochrome b5 (FC = 0.2, *p* = 3.85×10^-02^) exhibited the third-smallest FC value. It has been reported as a novel biomarker for visceral obesity intervention and a safe therapeutic target^104^, as well as a potential biomarker for distinguishing well-differentiated hepatocellular carcinoma^105^.

Tissue kallikrein (FC = 1.65, *p* = 1.77×10^-06^) exhibited the smallest *p*-value, and is involved in biological processes such as positive regulation of acute inflammatory response, positive regulation of apoptotic process, and regulation of systemic arterial blood pressure. It has been reported as a potential biomarker for renal papillary carcinoma and clear cell renal cell carcinoma^106^.

Acid ceramidase (FC = 1.96, *p* = 6.96×10^-05^) exhibited the third-smallest *p*-value. It possesses functions such as fatty acid amide hydrolase activity and N-acylsphingosine amidohydrolase activity. It is involved in biological processes such as cellular response to tumor necrosis factor, ceramide biosynthetic process, ceramide catabolic process, fatty acid metabolic process, regulation of programmed necrotic cell death, regulation of steroid biosynthetic process, and sphingosine biosynthetic process. *ASAH1* has been reported as a potential diagnostic biomarker for asthma^107^ and a plasma marker for the progression of recurrent glioblastoma^108^. ASAH1 can also serve as a biomarker for predicting lymph node status in patients with breast cancer^109^.

Lysosomal acid phosphatase (FC = 1.77, *p* = 4.62×10^-04^) exhibited the fifth-smallest *p*-value and has been reported as a biomarker for juvenile neuronal ceroid lipofuscinosis^110^.

Cadherin-13 (FC = 0.66, *p* = 7.23×10^-04^) has been reported to interfere with the differentiation potential of adipocytes, serving as a marker of adipose tissue plasticity and reflecting the health status of adipose tissue^111^. CDH13 can also serve as a prognostic biomarker and therapeutic target for renal clear cell carcinoma^112^.

##### 3.2.4.2 Analysis of Biological Pathways Enriched from Differential Proteins Identified by Comparison between the Pu-erh Tea Group and the Control Group

The differential proteins identified by comparison between the Pu-erh tea group and the control group were mainly involved in biological processes such as homophilic cell adhesion via plasma membrane adhesion molecules, inflammatory response, complement activation, cellular response to tumor necrosis factor, lipid metabolism, negative regulation of lipid storage, regulation of systemic arterial blood pressure, glycoside catabolism, hyaluronan metabolism, and mitral valve morphogenesis (Figure 10A).

**Figure 10.**
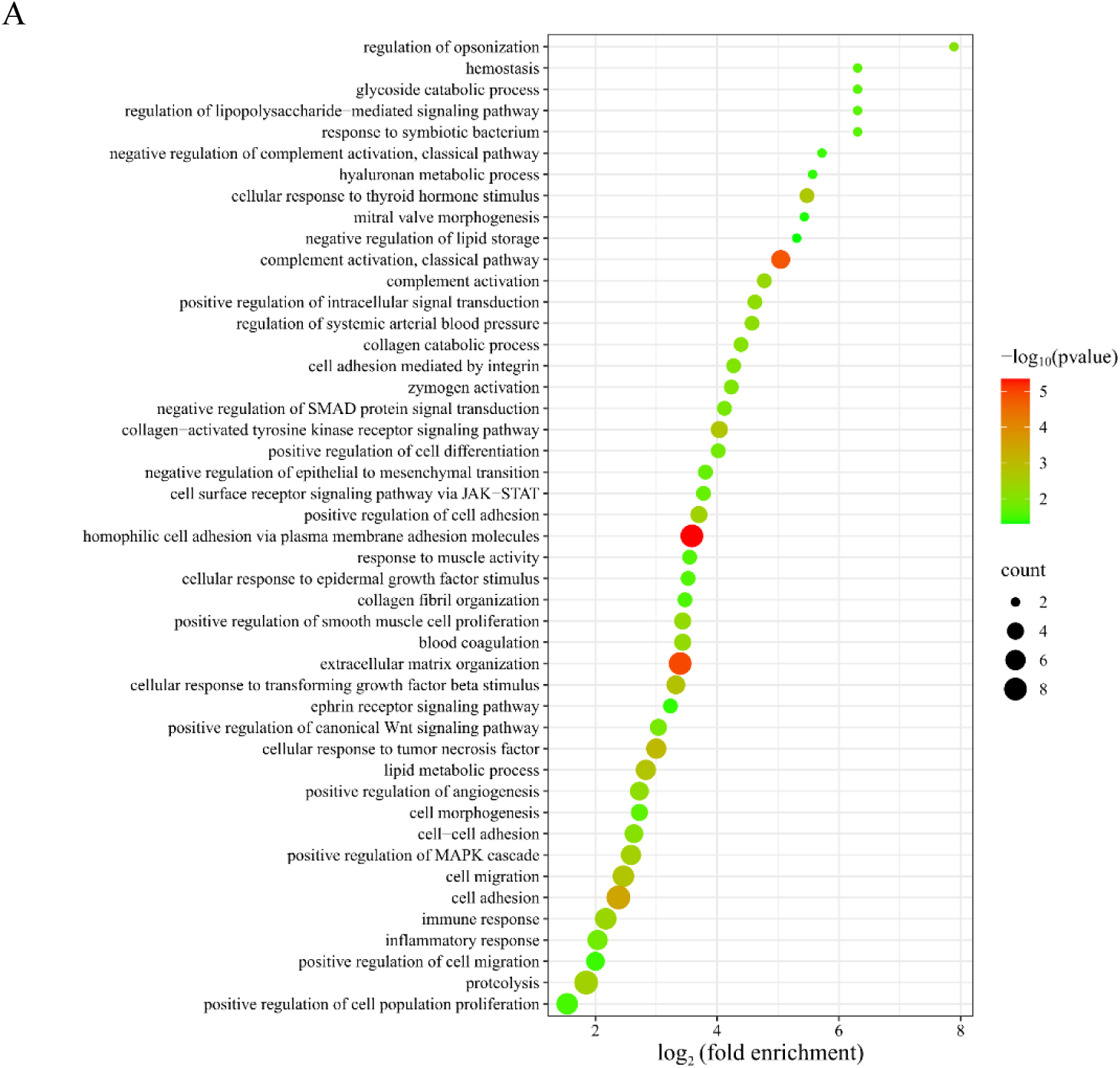
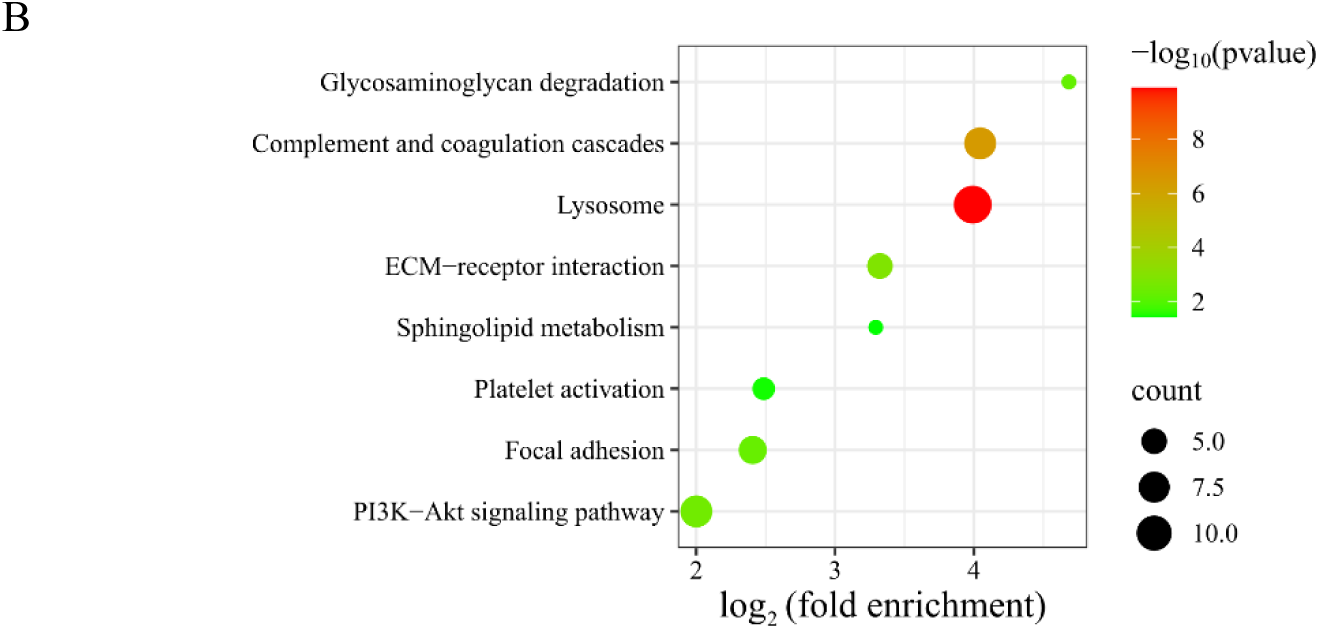
Functional annotation of differential proteins identified in the comparison between the Pu-erh tea group and the control group (*p* < 0.05): (A) biological processes; (B) KEGG pathways.

KEGG enrichment analysis revealed significant enrichment in pathways including lysosome, complement and coagulation cascades, ECM-receptor interaction, PI3K-Akt signaling pathway, glycosaminoglycan degradation, and sphingolipid metabolism (Figure 10B). Studies have shown that theaflavin, a bioactive compound in black tea, regulates glycolipid metabolism via the IRS-1/PI3K/Akt signaling pathway^46^. Additionally, Pu-erh tea extract has been reported to regulate alcohol-induced metabolic disorders via pathways such as sphingolipid metabolism^113^.

#### 3.2.5 Comparison Between the Black Coffee Group and the Control Group

By comparing the urinary proteins of rats in the black coffee group with those in the control group, a total of 46 differential proteins were identified under the screening conditions of FC ≥ 1.5 or ≤ 0.67 and *p* < 0.05 by two-tailed unpaired *t*-test analysis. Detailed information on these differential proteins is listed in Table S2. Results from the randomized grouping test showed that the average number of differential proteins yielded was 37.21, indicating that at least 19.11% of these differential proteins were not randomly generated.

##### 3.2.5.1 Analysis of Differential Proteins Identified by Comparison between the Black Coffee Group and the Control Group

Adhesion G protein-coupled receptor F5 (FC = 0.18, *p* = 4.47×10^-02^), exhibited the smallest FC value among the identified differential proteins, possesses the function of G protein-coupled receptor activity. It is involved in biological processes such as energy reserve metabolism, fat cell differentiation, glucose homeostasis, macrophage activation, negative regulation of macrophage activation, and phospholipid biosynthesis. ADGRF5 has been reported as a prognostic biomarker for renal clear cell carcinoma^114^. Plasma ADGRF5 levels have been reported as a potential biomarker for proliferative diabetic retinopathy^115^.

14-3-3 protein gamma (FC = 0.21, *p* = 1.23×10^-02^) exhibited the second-smallest FC value. It is involved in biological processes such as cellular response to insulin stimulus, negative regulation of TORC1 signaling, regulation of synaptic plasticity, and cellular response to glucose starvation. YWHAG has been reported as a potential prognostic biomarker and drug response biomarker for lung adenocarcinoma^116^, a potential prognostic biomarker and therapeutic target for oral squamous cell carcinoma^117^, a prognostic biomarker associated with shunt responsiveness in patients with idiopathic normal pressure hydrocephalus^118^, a potential cerebrospinal fluid biomarker for Alzheimer’s disease^119^, and a diagnostic biomarker for cognitive impairment in patients with Parkinson’s disease^120^.

ADAMTS-like protein 4 (FC = 0.23, *p* = 6.85×10^-03^) exhibited the third-smallest FC value and is involved in the apoptotic process. It has been reported as a biomarker for primary glioblastoma multiforme^121^ and a prognostic biomarker for Burkitt lymphoma^122^.

Enhancer of mRNA-decapping protein 4 (FC = 3.90, *p* = 2.99×10^-02^) exhibited the largest FC value and is involved in deadenylation-independent decapping of nuclear-transcribed mRNA and nervous system development.

Lymphocyte cytosolic protein 1 (FC = 2.92, *p* = 2.53×10^-02^) exhibited the second-largest FC value, possesses functions such as calcium ion binding, GTPase binding, and integrin binding. It is involved in biological processes including protein kinase A signaling and T cell activation involved in immune response. Studies have shown that LCP1 may serve as a potential therapeutic target for obesity-related metabolic disorders^123^, a biomarker for oral squamous cell carcinoma^124^, and a potential diagnostic and prognostic biomarker for triple-negative breast cancer^125^.

##### 3.2.5.2 Analysis of Biological Pathways Enriched from Differential Proteins Identified by Comparison between the Black Coffee Group and the Control Group

The differential proteins identified by comparison between the black coffee group and the control group were mainly involved in biological processes such as angiogenesis involved in wound healing, protein glycosylation, lysosome organization, regulation of lipopolysaccharide-mediated signaling pathway, lipid catabolic process, immune response, positive regulation of calcium ion import, and midbody abscission (Figure 11A). KEGG enrichment analysis revealed significant enrichment in pathways including lysosome, metabolic pathways, glycosphingolipid biosynthesis-lacto and neolacto series, sphingolipid metabolism, and glycosaminoglycan degradation (Figure 11B).

**Figure 11.**
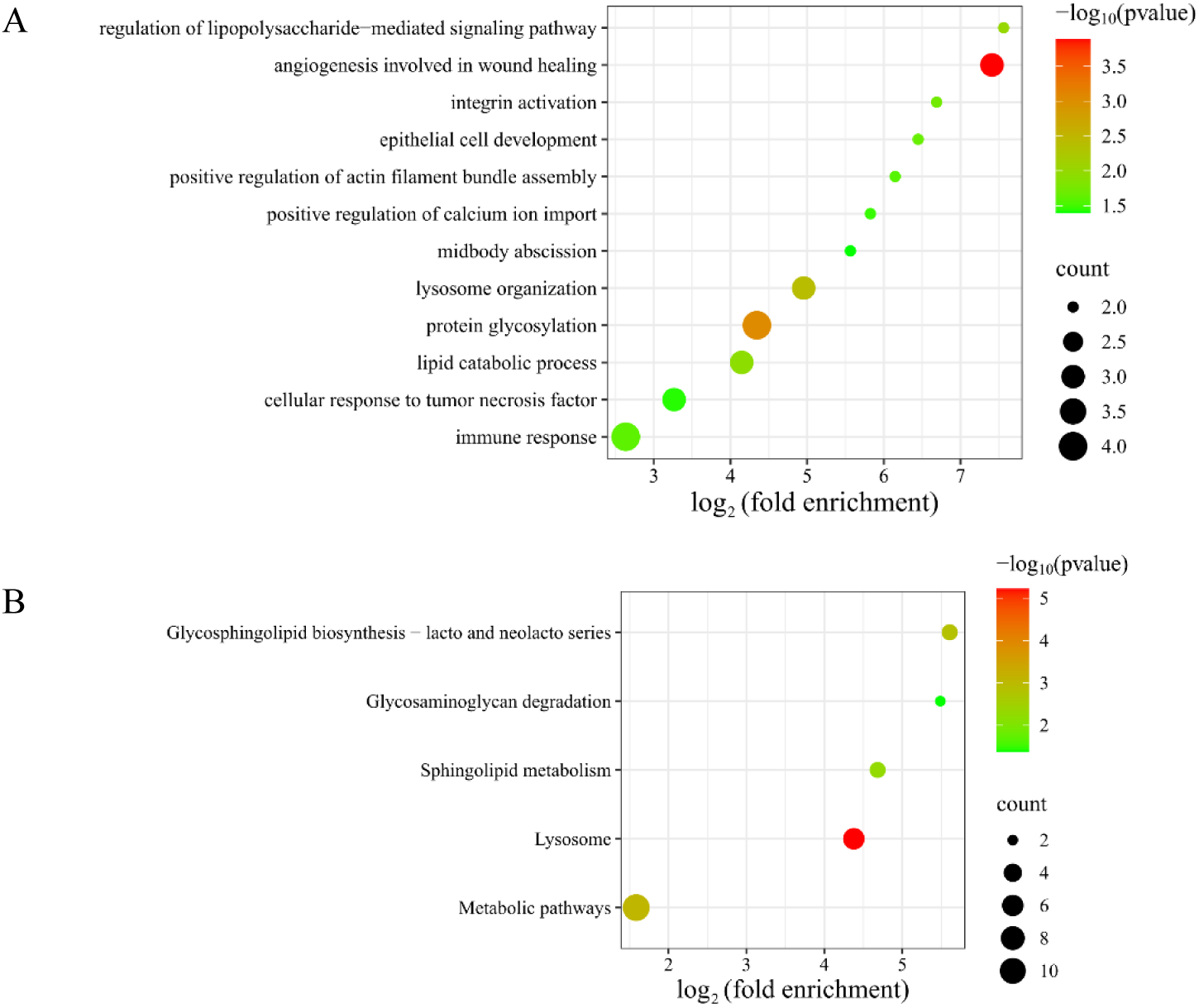
Functional annotation of differential proteins identified in the comparison between the black coffee group and the control group (*p* < 0.05): (A) biological processes; (B) KEGG pathways.

## 4 Discussion

The results of this study indicate that the urine proteome can reflect changes in rats after seven consecutive days of consuming teas with different fermentation levels and black coffee. The urinary proteins of rats in the control group (consuming sterile water) before and after 7 days were also compared in the experiment. Differential proteins identified in the control group were mainly involved in biological processes such as skeletal system morphogenesis, heart development, and cartilage development. These results indicate that the urine proteome can reflect short-term growth and development changes in rats, consistent with the previous results of our laboratory^126^. Therefore, to minimize the influence of various factors, before-and-after comparisons and between-group comparisons were performed for analysis. By minimizing the influence of individual differences through before-and-after comparisons and avoiding the effects of short-term growth and development in rats through between-group comparisons, the effects and differences of teas and black coffee were meticulously explored.

Venn diagrams were used to respectively show the overlap of differential proteins identified from the before-and-after comparisons and between-group comparisons in the four types of tea groups and black coffee group (Figure 12). The results showed that each group shared few common differential proteins, while many were unique to each group. Similarly, Venn diagrams of biological processes (Figure 13) and KEGG pathways (Figure 14) enriched from differential proteins across the five experimental groups revealed few shared elements and many unique ones among the groups. These findings suggest that urine proteome can distinguish the effects of teas with different fermentation levels and black coffee, and that these effects are highly specific to the types of beverage consumed.

**Figure 12.**
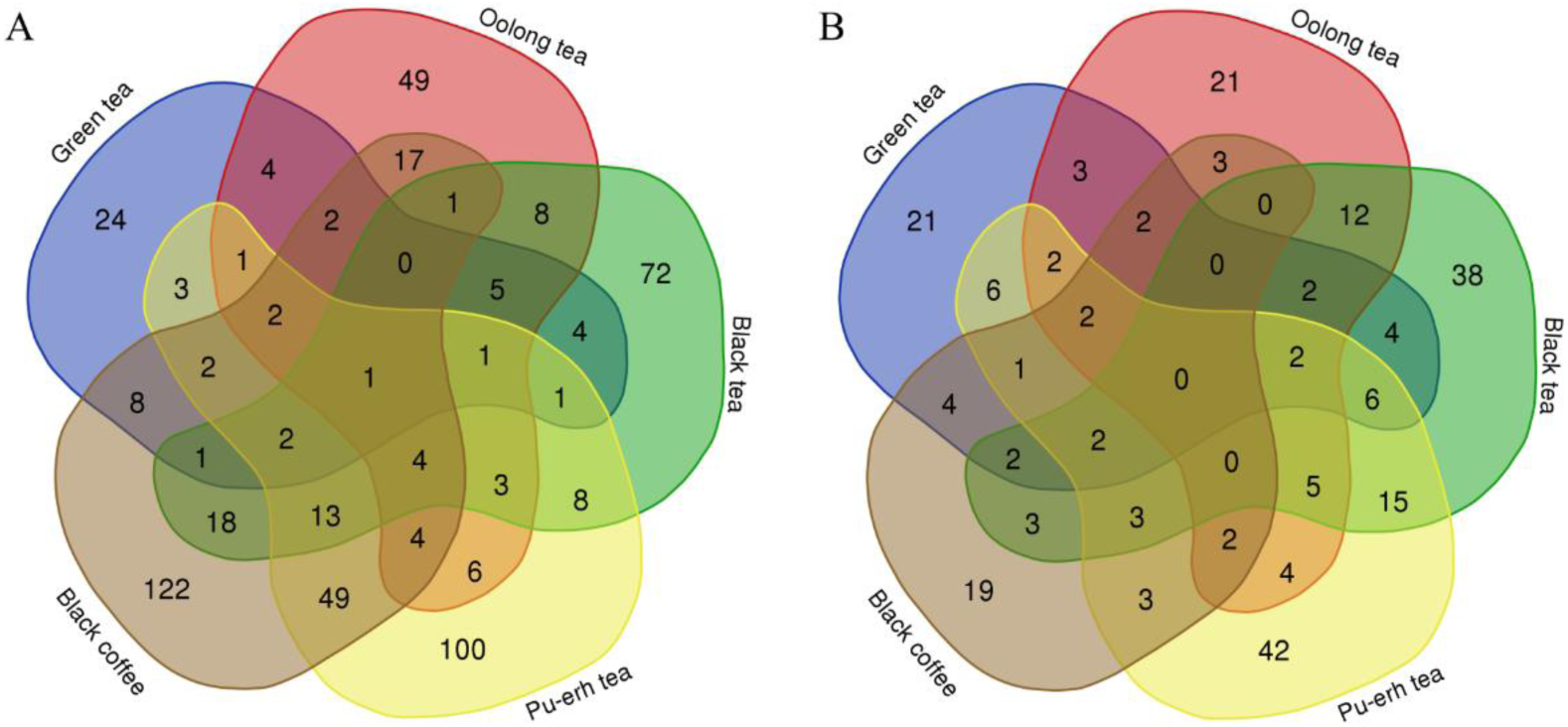
Venn diagrams of differential proteins across the five experimental groups: (A) before-and-after comparisons; (B) between-group comparisons.

**Figure 13.**
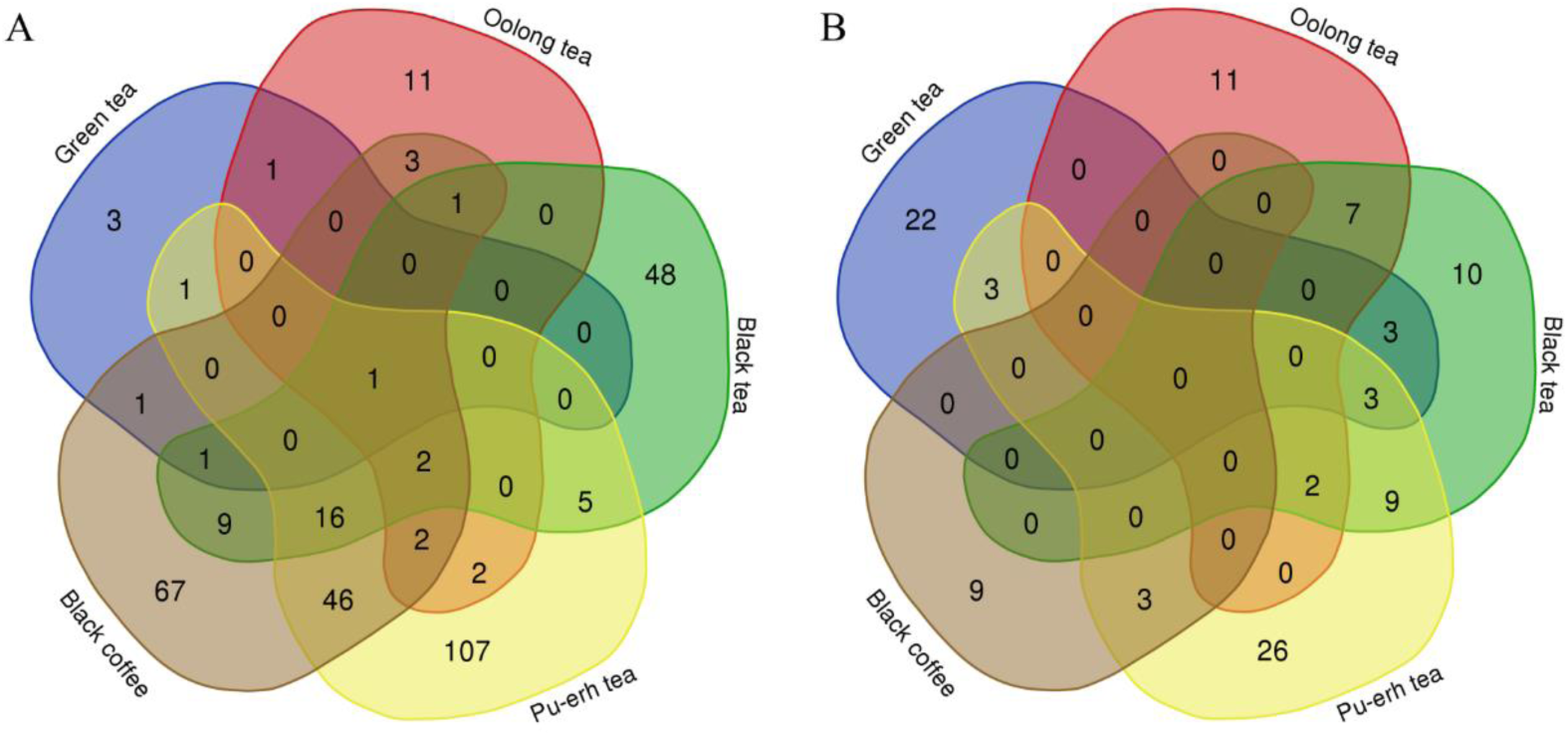
Venn diagrams of biological processes enriched from differential proteins across the five experimental groups: (A) before-and-after comparisons; (B) between-group comparisons.

**Figure 14.**
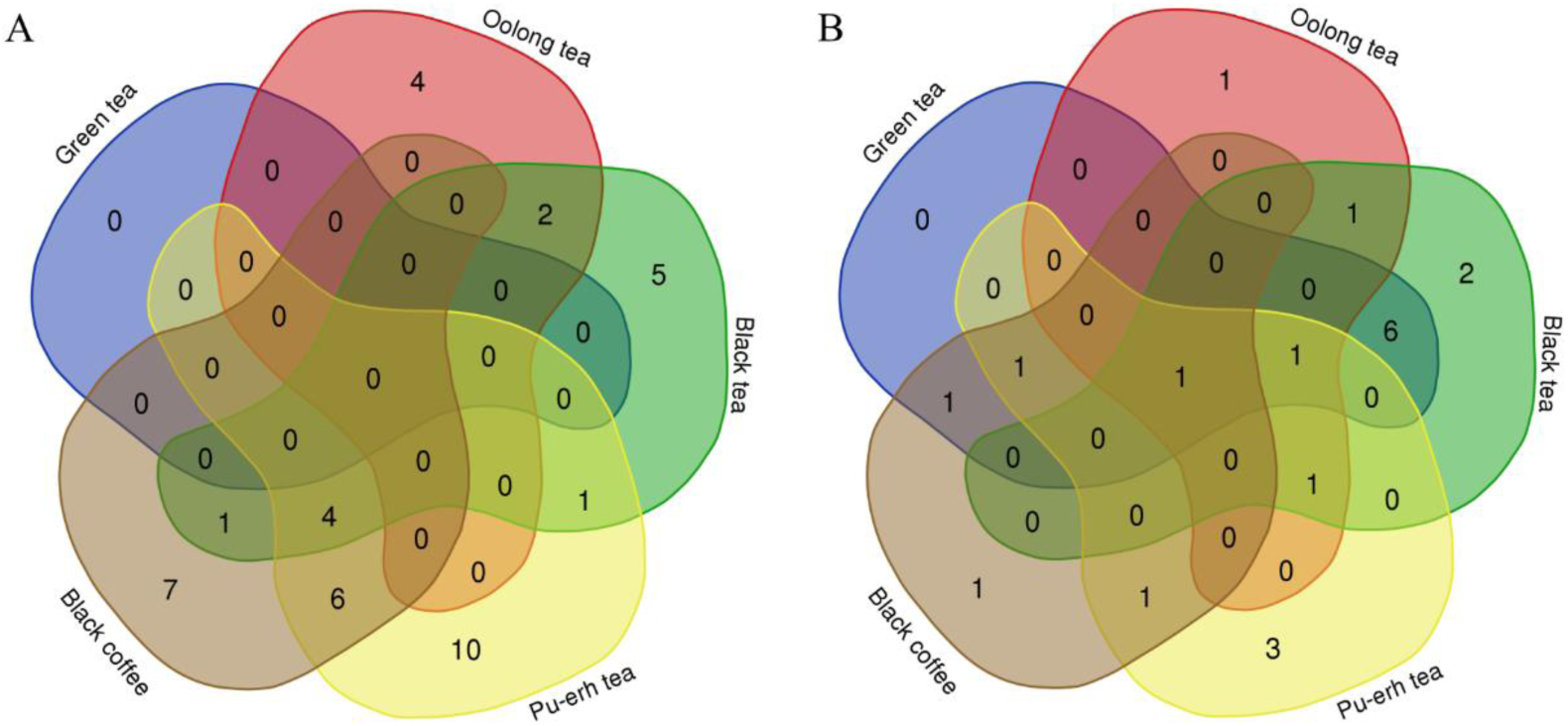
Venn diagrams of KEGG pathways enriched from differential proteins across the five experimental groups: (A) before-and-after comparisons; (B) between-group comparisons.

Additionally, this study found that multiple identified differential proteins have been reported as biomarkers for diseases such as cancer and cardiovascular diseases, particularly those proteins that exhibit significant changes under the influence of teas and black coffee. This provides references for the research of urine biomarkers of diseases and related studies, suggesting that future research and clinical applications of urine biomarkers of diseases should consider the potential influence of beverage consumption, such as tea and black coffee. During clinical urine sample collection, it is necessary to consider whether beverage restrictions are required. Additionally, relying on a single biomarker may lead to false positives, so it may be necessary to use biomarker panels to improve accuracy.

## 5 Conclusion

The urine proteome comprehensively and systematically reflects changes in rats after seven consecutive days of consuming four types of teas or black coffee, and distinguishes the changes in the body after consuming teas with different fermentation levels and black coffee.

